# *Neisseria gonorrhoeae* subverts formin-dependent actin polymerization to colonize human macrophages

**DOI:** 10.1101/2021.01.19.427361

**Authors:** Stanimir S. Ivanov, Reneau Castore, Magdalena Circu, Ana-Maria Dragoi

## Abstract

Dynamic reorganization of the actin cytoskeleton dictates plasma membrane morphology and is frequently subverted by bacterial pathogens for entry and colonization of host cells. The human-adapted bacterial pathogen *Neisseria gonorrhoeae* can colonize and replicate when cultured with human macrophages, however the basic understanding of how this process occurs is incomplete. *N. gonorrhoeae* is the etiological agent of the sexually transmitted disease gonorrhea and tissue resident macrophages are present in the urogenital mucosa which is colonized by the bacteria. We uncovered that when gonococci colonize macrophages they can establish an intracellular or a cell surface-associated niche that support bacterial replication independently. Unlike other intracellular bacterial pathogens, which enter host cells as single bacterium, establish an intracellular niche and then replicate, gonococci invade human macrophages as a colony. Individual diplococci are rapidly phagocytosed by macrophages and transported to lysosomes for degradation. However, we found that surface-associated gonococcal colonies of various sizes can invade macrophages by triggering actin skeleton rearrangement resulting in plasma membrane invaginations that slowly engulf the colony. The resulting intracellular membrane-bound organelle supports robust bacterial replication. The gonococci-occupied vacuoles evaded fusion with the endosomal compartment and were enveloped by a network of actin filaments. We demonstrate that gonococcal colonies invade macrophages via a process mechanistically distinct from phagocytosis that is regulated by the actin nucleating factor FMNL3. Our work provides insights into the gonococci life-cycle in association with human macrophages and defines key host determinants for macrophage colonization.

## INTRODUCTION

The betaproteobacteria *Neisseria gonorrhoeae (N*.*g)* is a highly adapted human colonizer and the etiological agent of the sexually transmitted disease gonorrhea. Recently, gonorrhea has emerged once again as a major global public health problem due to increased incidence as well as rapid emergence of antibiotics resistance (1, 2). Majority of gonococcal infections are asymptomatic and human-to-human transmission maintains the gonococcal reservoir within the human population (3, 4). Gonococci colonize the urogenital tract, often trigger a localized inflammatory response, which in acute symptomatic infections can progress to the production of purulent exudate that contains gonococci, innate immune cells (macrophages and polymorphonuclear leukocytes-PMNs) and exfoliated epithelial cells (5, 6). A disseminated gonococcal infection can lead to pelvic inflammatory disease, scarring of the fallopian tubes, arthritis and endocarditis (7-9).

Depending on the exposure route in human infections, gonococci have been found to colonize several mucosa including the genital, ocular, nasopharyngeal and anal (8). In tissues, gonococci adhere to the host epithelium, proliferate, invade the submucosal region and encounter tissue-resident innate immune cells (3). Robust immune evasion mechanisms directed towards both the innate and the adaptive immune system allows gonococci to establish persistent infections (10-19). For example, high frequency phase and antigenic variation diversifies cell surface exposed polypeptides to counteract host adaptive immunity (20-23). Also, gonococci colonize and replicate in association with immune phagocytes including macrophages and neutrophils (10-12, 24-27). The molecular determinants that dictate epithelial cells and neutrophils colonization are well understood; however, macrophage colonization by gonococci remains enigmatic. Macrophages could potentially support gonococcal replication during infection as gonococci survive and replicate when cultured with human primary and cell line-derived macrophages (10, 28, 29). Macrophages represent ∼10% of total leukocytes isolated from the genitourinary mucosa (5) and are recovered from sites of acute gonococcal infections (6). *N*.*g* blocks staurosporine-induced apoptosis in human primary and monocytic cell line-derived macrophages while eliciting both inflammatory and immune suppressive cytokines (10, 13). In macrophage infections, the origin and location of the niche supporting gonococcal replication remains an important open question as gonococci reside within lysosomes (10, 30) in the host cytosol (10, 31, 32) and on the cell surface (10, 32, 33).

Gonococci tether to host cells via type IV pili (34-37) and produce a number of adhesins that engage surface receptors to facilitate tight association with epithelial cells and neutrophils (3). The outer membrane protein PorB and the Opacity (Opa) proteins family are key gonococcal adhesins (38-43). In general, gonococcal strains encode up to 11 *opa* loci (44, 45) which give rise to 7 to 9 unique Opa proteins that bind to the human carcinoembryonic antigen-related cell adhesion molecule (CEACAM) surface receptor family (46). Different Opa proteins preferentially bind distinct CEACAM receptors and interactions with CEACAM1, CEACAM3, CEACAM5 and CEACAM6 have been reported (47-52). Opa binding to the broadly expressed CEACAM1 mediates attachment to neutrophils and epithelial cells (52), whereas engagement of the neutrophil specific CEACAM3 receptor transports gonococci to a degradative compartment (53-55). The *opa* alleles undergo high frequency phase variation, which diversify the combination of Opa proteins produced by each diplococcus within a bacterial colony (46). Thus, expression of the Opa adhesins benefit host cell colonization at the cost of enhanced killing by neutrophils.

Gonococci and the closely related meningococci are considered epicellular pathogens (56) that form large colonies on the surface of the host cell where plasma membrane proximal bacteria are embedded within membrane ruffles enriched in polymerized actin microfilaments, also known as actin plaques (57-61). A number of host cell surface receptors and adhesion molecules (EGFR, CD46, CD44 and ICAM) that engage the cortical actin network aggregate within the gonococci-induced actin plaques (60, 62). Ezrin – an adaptor protein that link surface receptors with the cortical actin cytoskeleton is also enriched within these structures (57, 60). Actin plaques are likely produced as a result of surface receptor clustering by the gonococci type IV pili as formation depends on the cholesterol content of the plasma membrane (63). Internalized diplococci have been observed in epithelial cells; however, they represent a minor fraction of the total cell-associated bacterial population (60), thus whether and to what extent intracellular bacteria contribute to overall gonococcal expansion remain unclear. Epicellularity is thought to confer some protection from bactericidal host factors especially of the membrane embedded fraction of the bacterial colony (56).

Dynamic remodeling of the actin cytoskeleton by bacterial pathogens is a critical mechanism for host invasion and spreading (82, 83). Both Arp2/3-dependent and formin-dependent mechanisms are hijacked by bacterial pathogens in order to polymerize actin and enable cell entry and cell-to-cell spreading. While Arp2/3 complex initiates branched actin formation by binding to the sides of pre-existing filaments, formin family of proteins directs the elongation of unbranched actin filaments.

Here, we investigate the molecular processes mediating human macrophage colonization by gonococci and define distinct subcellular niches occupied by the bacteria that support replication. Also, we provide evidence for a novel mechanism that allows bacterial colonies rather than individual diplococci to invade and colonize human macrophages intracellularly. We demonstrate that the formin actin nucleator FMNL3 localizes at the sites of invasion and is required for *N*.*g* colony internalization. These results suggest that *N*.*g* subverts FMNL3 functions in actin cytoskeleton remodeling to facilitate colonization of human macrophages.

## Results

### N.g colonizes and replicates in association with human macrophages

We investigated the capacity of *N*.*g* to replicate in association with human macrophages by quantifying the bacterial colony forming units (CFUs) recovered at discrete times after infection. In these infections, we used PBSG (PBS, 7.5mM glucose, 900µM CaCl_2_, 500µM MgCl_2_) because this media by itself does not support gonococcal replication (10) (Fig. 1a). However, when human macrophages differentiated from the pro-monocytic U937 cell line were cultured with different *N*.*g* strains in PBSG, the gonococcal CFUs increased by 2 to 3 log_10_ in an 8-hour period (Fig. 1a-b). Under those conditions, the percentage of infected cells increased from 2 to 28% and the average number of gonococci per macrophage also increased from 1.6 to 47.7 (Fig. 1c-e and Fig. S1). The growth kinetics in macrophage infections for strains isolated from patients with disseminated (FA1090, FA19) or uncomplicated (F62, MS11) gonococcal infections (64) were comparable and exceeded the *in vitro* growth rate in the complex gonococcal base liquid (GCBL) medium (Fig. 1b). Taken together, these data from cell culture infections with a low starting multiplicity of infection (MOI) demonstrate gonococci capacity to colonize human macrophages, replicate and disseminate to neighboring bystander cells within hours.

**Figure 1.**
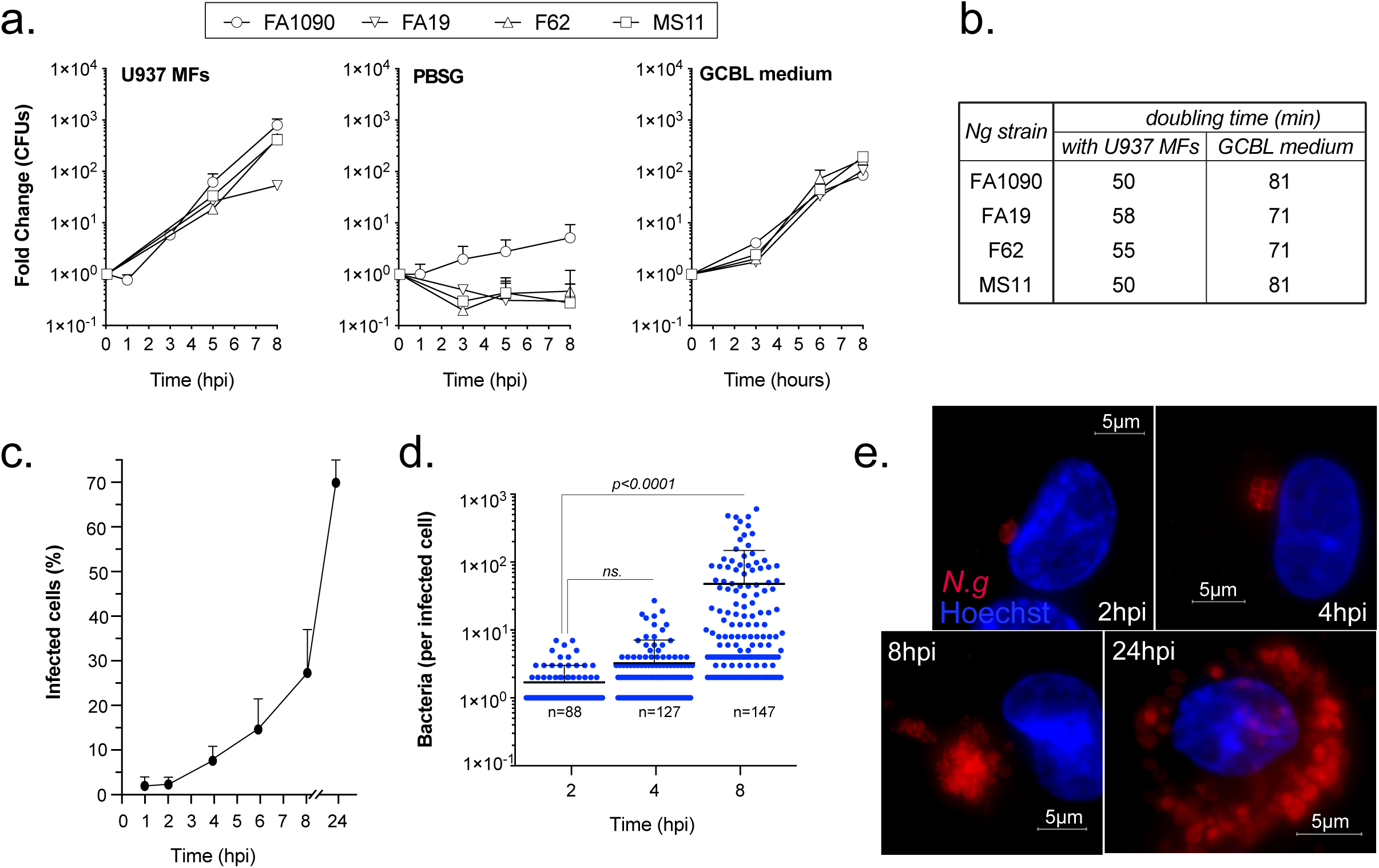
*N. gonorrhoeae* associates and replicates with human macrophages. **(a)** Growth of different *N*.*g* strains cultured in the presence/absence of differentiated U937 human macrophages (U937 MFs) at MOI=0.1 or in axenic rich liquid culture media GCBL. **(b)** Growth rate of gonococcal strains when cultured with U937 MFs or in GCBL media calculated from recovered CFUs counts. **(c-e)** Microscopy analysis of gonococcal growth kinetics of *N*.*g* FA1090 with U937 MFs at MOI=0.1. The percentage of infected macrophages **(c)** and the number of bacteria per cell **(d)** are shown. **(e)** Representative micrographs of *N*.*g* FA1090 infected cells at the indicated time-points. **(a** and **c)** Means of technical replicates ± SD are shown. **(d)** Each point represents a single macrophage; means with SD are shown; unpaired T-test analysis, not significant (ns.)

When gonococci were separated from the U937 macrophages (U937 MFs) with a 0.4µm filter barrier that allowed fluid exchange but prevented direct contact, gonococci replicated poorly (Fig. 2a). In addition, macrophages fixed with paraformaldehyde prior to infection also did not support gonococcal replication (Fig. 2b), indicating that direct contact with live macrophages is required to stimulate robust bacterial replication. Together, these data demonstrate that gonococci likely have evolved mechanisms to colonize and replicate in association with human macrophages.

**Figure 2.**
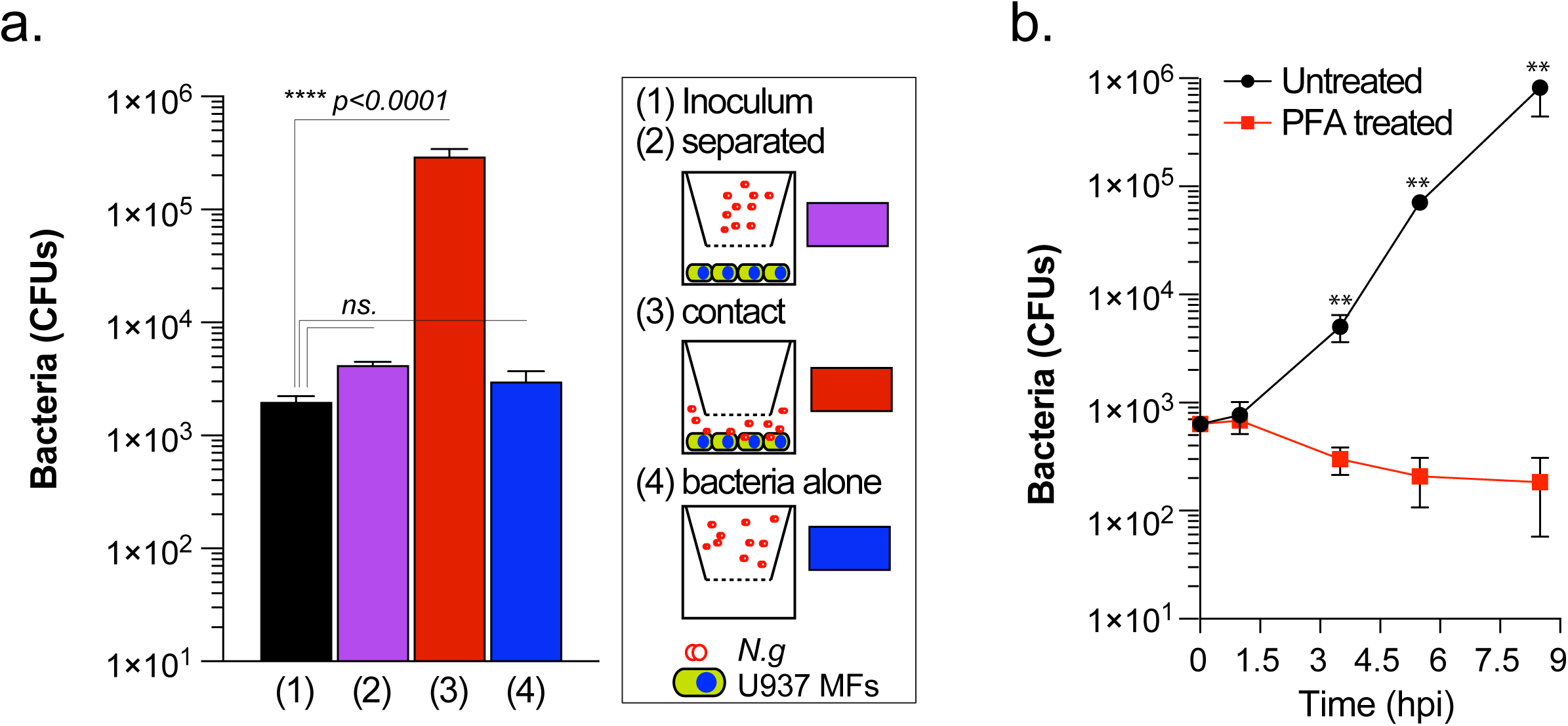
Direct contact with live macrophages is required for gonococcal replication. **(a)** *N*.*g* FA1090 growth under conditions that permit or restrict direct contact with U937 MFs in PBSG media. Contact separation between bacteria and the macrophages is facilitated by a 0.4µm filter barrier. Total CFUs recovered at 8hpi are shown. **(b)** *N*.*g* FA1090 growth in association with PFA-fixed macrophages in PBSG media. Macrophages were fixed for 45min with PFA (4% vol./vol. in PBS) and extensively washed with warm PBS prior to infection. Total CFUs recovered at the indicated time-points are shown. **(a-b)** Means ± SD are shown, ** p < 0.005 unpaired T-test.

### N.g occupies multiple distinct cellular niches on human macrophages

Because macrophages are phagocytes, it is not surprising that intracellular as well as cell surface-associated gonococci have been observed in previous studies (10); however, the subcellular niche that supports gonococcal replication remains unknown. Thus, we first investigated where gonococcal colonies were found on macrophages, because those subcellular compartments likely support bacterial replication. To this end, 3D immunofluorescence microscopy analysis of U937 MFs infected with *N*.*g* FA1090 for 6 hours was performed, where the cortical actin network was labeled with phalloidin (Fig. 3a). Cortical actin can discern intracellular from cell-surface localized bacteria as it is formed in close proximity to the plasma membrane. Based on this criterion, both intracellular and cell surface-associated gonococcal colonies were observed (Fig. 3a). These distinct colony topologies were confirmed by an alternative inside/out immunofluorescence microscopy approach, in which extracellular bacteria are labeled with two distinct polyclonal anti-*N*.*g* antibodies, one used prior to and another used after cell permeabilization, whereas intracellular bacteria are single-labeled because they are accessible only after cell permeabilization (Fig. 3b). Surprisingly, multiple colonies had mosaic inside/out staining pattern consisting of single antibody-stained as well as dual-stained regions within the same colony (Fig. 3c). This unique hybrid colony type was produced by all four strains tested (FA1090, MS11, F62, and FA19) in U937 MFs infections (Fig. 3d) as well as in infections of human primary monocyte-derived macrophages (hMDMs) (Fig. 3e and Fig. S2c-d). These data demonstrate that macrophage colonization produces gonococcal colonies with distinct topologies by a mechanism that is broadly conserved in primary hMDMs and U937 MFs.

**Figure 3.**
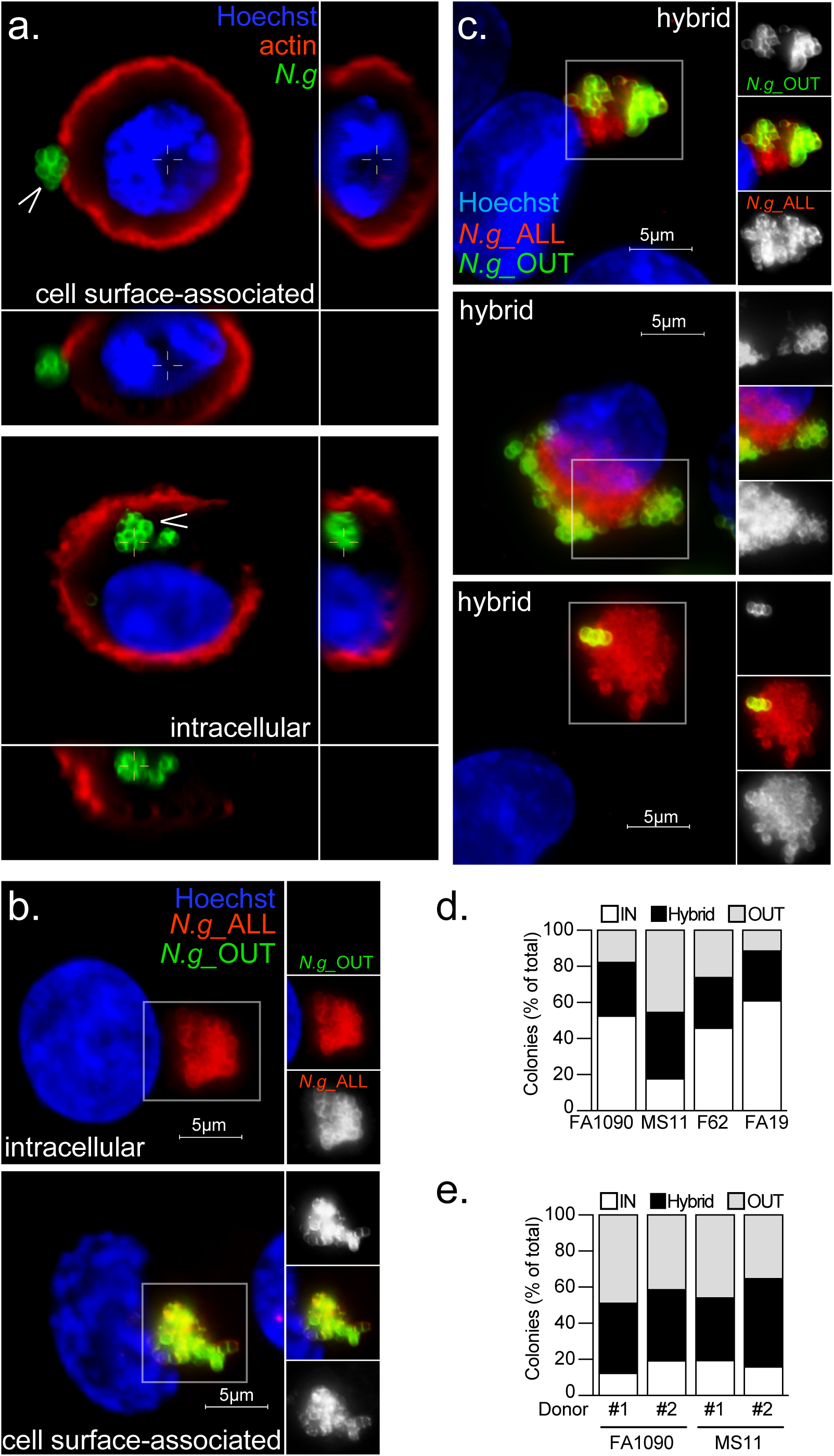
Gonococci colonize distinct cellular compartments. **(a-c)** U937 MFs infections with *N*.*g* FA1090 at MOI=2. **(a)** Representative single focal plane of 3D image Z-stacks showing intracellular and cell-surface associated gonococcal colonies at 6 hpi. Side panels show Z-axis cross-sections at the indicated areas. **(b-c)** micrographs of colonized macrophages after inside/out staining with anti-*N*.*g* antibodies. **(b)** Single-stained intracellular (red) and double-stained surface-exposed (yellow) gonococcal colonies are shown. **(c)** Multiple representative partially exposed gonococcal colonies (yellow and red) are shown. **(d-e)** Quantitative analyses of the relative colony topologies distribution at 10 hpi in U937 MFs **(d)** and in human primary monocyte-derived macrophages (hMDMs) **(e)** are shown.

We considered that the hybrid colony topology could be a type of surface-associated colony in which tightly packed outer layer gonococci render innermost bacteria inaccessible to antibody binding. However, 3D microscopy clearly demonstrates that at sites where colonies with hybrid topology are present a meshwork of cortical actin envelops the cell proximal regions of the gonococcal colony rendering those colony regions inaccessible to the antibodies (Fig. 4a). Thus, in the hybrid colonies, single antibody stained regions are likely intracellular and the dual stained regions are surface exposed. Although the intra-and extracellular regions in hybrid colonies are clearly distinguishable by inside/out staining, hybrid colonies remain continuous and appear as a single entity (Fig. 4b), indicating that the bacteria likely occupy a unique quasi-intracellular compartment where the extracellular gonococci acts as a plug to seal a plasma membrane invagination harboring the intracellular portion of the colony. Indeed, plasma membrane invaginations that co-localize with the actin meshwork at the cell-proximal regions of hybrid gonococcal colonies are evident (Fig. 4c). Together, these data indicate that in human macrophages gonococcal colonies are found in three discrete locations: (1) on the cell surface, (2) within an intracellular compartment and (3) in a hybrid quasi-intracellular compartment.

**Figure 4.**
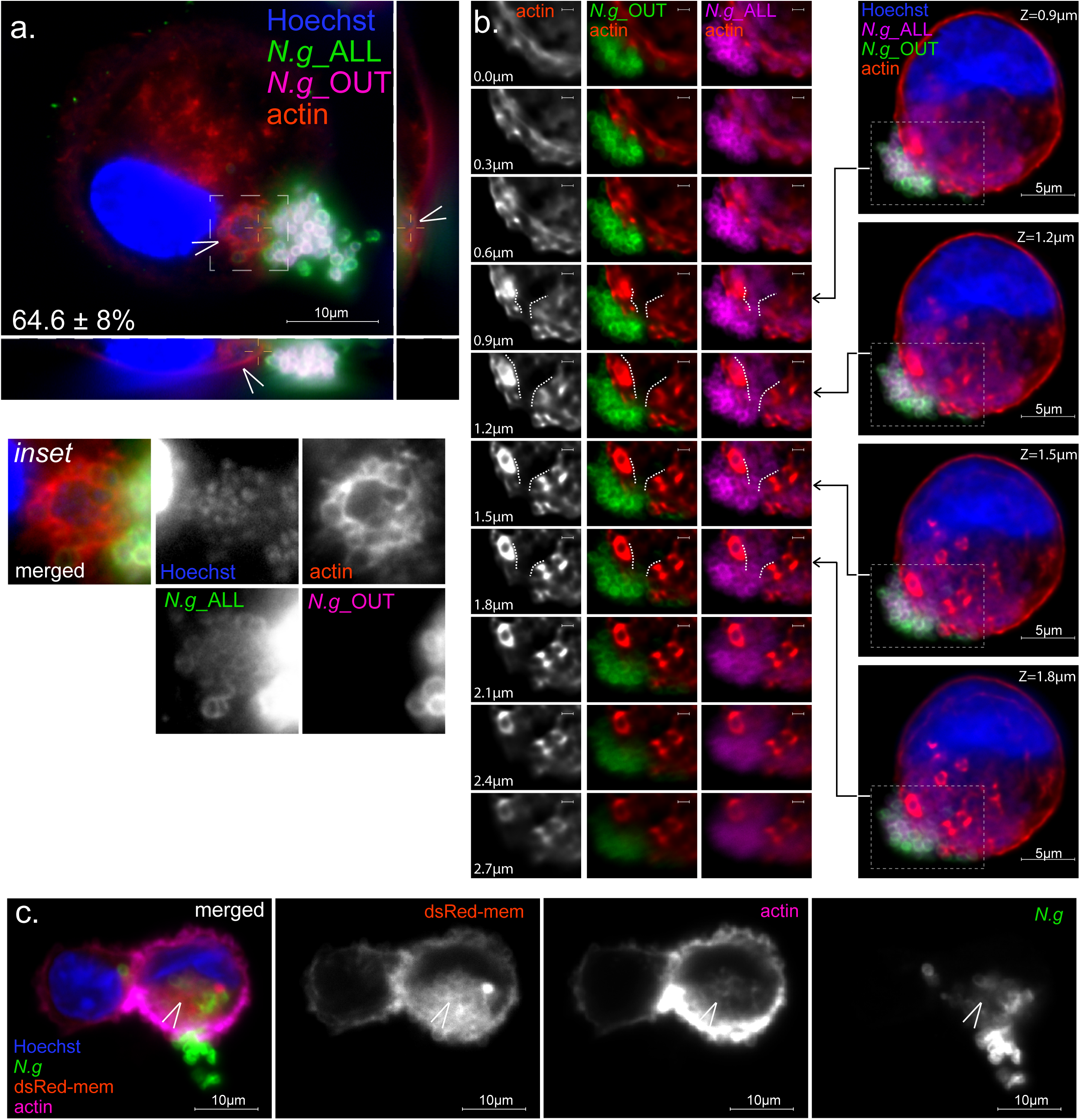
Actin microfilaments and membrane invaginations localize at plasma membrane sites colonized by hybrid gonococcal colonies. Micrographs show representative embedded gonococcal colonies on U937 MFs (MOI=2) at 8hpi **(a)** and 10hpi **(b)** after inside/out differential staining. **(a)** Side panels show Z-axis reconstruction of the indicated cross-section. Inset shows polymerized actin accumulation (found on 64.6 ± 8% of embedded colonies, n=54) specifically on the intracellular region of the hybrid colony (arrowhead). **(b)** Embedded colonies are continuous. A series of single focal planes spanning 2.7µm shows the transition point-from intracellular to extracellular region of an embedded colony is continuous and lined by cortical actin. **(c)** Actin-rich plasma membrane invaginations accumulate on the intracellular region of hybrid colonies (arrowhead). Single focal plane micrograph of a U937 MF stably expressing the plasma membrane-associated fluorescent reporter DsRed-mem and harboring a hybrid gonococcal colony.

### Intracellular gonococcal colonies arise from prolonged internalization of surface-associated gonococcal microcolonies

In the first hour of infection, over 40% of live as well as heat-killed gonococci were internalized by U937 MFs (Fig. S3a) The majority of those bacteria (both live and heat-killed) localized in lysosomes (Fig. 6d), indicating that rapidly internalized diplococci could be transported to lysosomes for degradation. Indeed, pre-treatment with lysosome inhibitors significantly increase the number of surviving intracellular gonococci in U937 MFs infections (10). If rapidly phagocytosed diplococci are likely eliminated, we sought to determine when productive entry events that give rise to intracellular colonies occur. To this end, we blocked phagocytosis by adding cytochalasin D at discrete times post infection to prevent bacterial uptake and scored the number of intracellular colonies formed by 8 hpi (Fig. 5a). As expected, pre-treatment with cytochalasin D blocked the biogenesis of intracellular colonies. However, blocking phagocytosis as late as 3 hpi reduced the total number of intracellular and hybrid colonies detected at 8hpi by 79% (from 80% to 17%). These results further support the idea that rapidly phagocytosed diplococci do not give rise to intracellular colonies and demonstrate that productive intracellular colonization is likely initiated by bacteria that are internalized at or later then 3hpi. Similar results were obtained when a gentamicin (a membrane impermeable antibiotic) protection assays was used (Fig. 5b), which showed complete sensitivity to the drug at 1 hpi and a significant increase in the protected population from 2 to 4 hpi. Thus, intracellular colonization is carried out by gonococci that remain surface-associated for at least three hours post adherence, whereas the majority of early entry events (< 3hpi) failed to establish intracellular infection.

**Figure 5.**
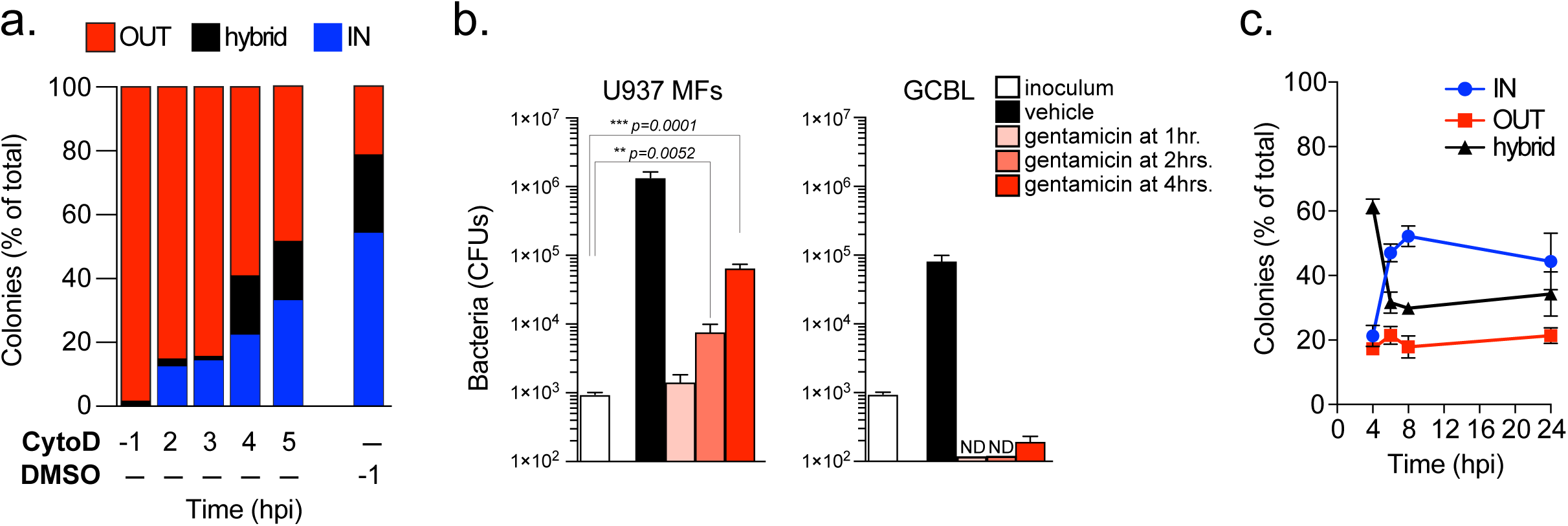
Kinetics of intracellular colonization by gonococci. **(a)** Late entry facilitates intracellular colonization. Break down of colony types found on U937 MFs infected with *N*.*g* FA1090 at 8hpi when the actin polymerization inhibitor cytochalasin D (5 µM) was added at the indicated time points to block bacterial internalization. Bacterial internalization is significantly blocked even when cytochalasin D is added as late as 3hpi. At least 100 colonies for each condition were scored. **(b)** *N*.*g* survives intracellularly in U937 MFs upon late gentamicin treatment. Bacterial CFUs recovered at 8hrs following a gentamicin protection assay in infected U937 MFs or in liquid culture (GCBL) are shown. Gentamicin (4µg/ml) was added at the indicated time-points. **(c)** Kinetic analysis of colony types in U937 MFs infections (MOI=2) with *N*.*g* FA1090. **(b-c)** Means of technical triplicates ± SD, unpaired T-test.

**Figure 6.**
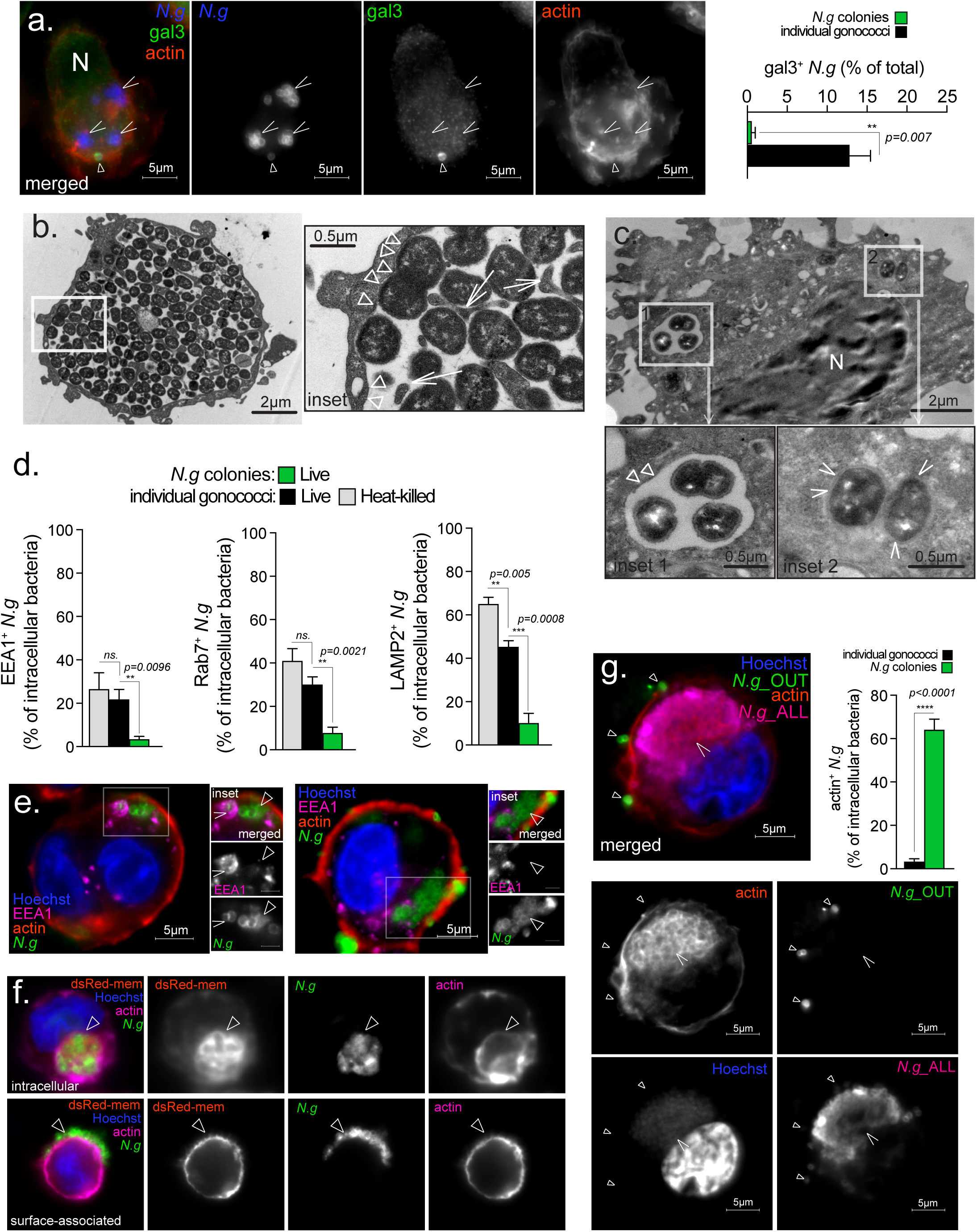
Gonococcal colonies evade endocytic maturation upon entry in macrophages and occupy a unique membrane-bound organelle surrounded by actin filaments. **(a)** Representative single focal plane micrograph showing Galectin-3 recruitment to some intracellular gonococcus (triangle) but not to gonococcal colonies (arrowhead). Quantitative analysis of galectin-3 recruitment to intracellular gonococci at 4hpi. Means of technical triplicates ± SD are shown, unpaired T-test. At least 100 objects for each condition were scored. **(b-c)** Representative TEM micrographs showing large **(b)** and small **(c)** intracellular gonococcal colonies that are separated from the cytoplasm by a membrane (insets, triangles) **(b-c)**, whereas some individual gonococcus can be seen in intimate contact with the cytoplasm (inset, arrowheads). **(b)** Membrane invaginations within the lumen of the *N*.*g*-occupied vacuole in close contact with gonococci were seen (inset, arrows). **(d-e)** Endosomal maturation microscopy analysis of *N*.*g* FA1090-occupied vacuoles. **(d)** Quantitative analysis of Early Endosome Antigen 1 (EEA1) and late endosomal markers Rab7 and LAMP2 recruitment to vacuoles occupied by heat-killed or live individual gonococci at 2hpi as well as internalized gonococcal colonies at 6hpi. At least 100 objects were scored for each condition. Means of technical triplicates ± SD are shown, unpaired T-test. **(e)** Representative single focal plane micrograph showing EEA1 accumulation on a vacuole containing a diplococci (arrowhead) and exclusion from the compartment occupied by *N*.*g* colony (triangles). **(f)** The membrane-bound compartment occupied by *N*.*g* colonies retains the plasma membrane marker DsRed-mem. Representative single focal plane micrographs showing internalized and surface-associated colonies with accumulation of DsRed-mem surrounding the intracellular bacterial colony. **(g)** Polymerized actin filaments accumulate on intracellular *N*.*g* colonies. Quantitative analysis and a representative single focal plane micrograph showing actin accumulation on an intracellular colony (arrowhead) at 10 hpi. Triangles point to surface-associated diplococci. Individual channels of the merged image are shown in grayscale. Infections of U937 macrophages with *N*.*g* FA1090 are shown. At least 85 objects were scored for each condition. Means of technical triplicates ± SD are shown, unpaired T-test.

To identify when and where gonococci initiate replication we investigated the appearance and cellular distribution of gonococcal microcolonies (4 – 12 bacteria) as a readout for initial bacterial replication. At 2 hpi microcolonies were already detectable when macrophages were infected with live but not heat-killed gonococci demonstrating that microcolonies result from bacterial replication rather than aggregation (Fig. S3b). Consistent with early replication by a few adherent gonococci the majority of bacteria were diplococci (∼ 90%) at that time. At 2hpi, a minor fraction of microcolonies were intracellular (< 13%) whereas the majority had hybrid topology (> 73%) (Fig. S3c). At 4hpi, the percentage of intracellular microcolonies increased and number of hybrid colonies decreased demonstrating that gonococci replicate first at the cell surface for at least 1-2 rounds to produce microcolonies, which are subsequently internalized to become intracellular. In agreement, colonies with hybrid topology were the most prevalent colony type at 4 hpi (∼ 60% of all colonies) but gradually declined by 50% at 8hpi (Fig. 5c). Conversely, intracellular colonies had the opposite trajectory and sharply increased in the same time period (Fig. 5c) providing evidence that hybrid colonies represent a snapshot of the internalization process.

The percentage of surface-associated extracellular colonies remained constant over a 24hrs period (Fig. 5c) indicating that surface-associated gonococci likely replicate to form colonies. We hypothesize that intracellular colonies originate from these surface-associated colonies after internalization. Because of the high phagocytic rate, persistent surface colonization by *N*.*g* in macrophages is surprising given that *N*.*g*-colonized macrophages internalized opsonized (IgG-coated beads) as well as non-opsonized (*L. pneumophila ΔdotA* bacteria) cargo normally (Fig. S4).

Indeed, large surface-associated colonies were detected as the infection progressed (Fig. 6f). Gonococcal rate of replication on macrophages was not reduced in the presence of cytochalasin D (Fig. S5a-b), providing further support that gonococci replication can occur independent of internalization.

Remarkably, once gonococcal colonies are internalized, they continued to expand even after bacterial uptake was blocked with cytochalasin D at rates comparable to the growth of surface-associated colonies and intracellular colonies in the absence of cytochalasin D (Fig. S5c-d). This can only occur if gonococci continue to replicate intracellularly. Thus, gonococci have the capacity to establish an intracellular as well as a cell surface-associated niche that support bacterial replication independently in macrophage infections.

### A unique actin-enriched membrane-bound organelle supports the replication of intracellular gonococci in macrophage infections

Although it has been speculated that gonococci escape in the cytosol of macrophages to replicate (10), direct evidence for cytosolic replication is lacking and thus the origin and identity of the intracellular niche supporting gonococci replication is still unclear. First, we sought evidence for cytosolic replication. The cytosolic β-galactoside-binding protein galectin-3 is frequently used as a marker to identify bacteria that invade the host cytosol because it binds sugar moieties present on the bacterial surface as well as on luminal glucosylated proteins that become exposed upon rupture of vacuolar compartments (72, 73). Within macrophages, a minor percentage of individually dispersed diplococci accumulated galectin-3, whereas colonies remained inaccessible to galectin-3 (Fig. 6a). Moreover, transmission electron microscopy analysis of infected internalized gonococci confirmed that colonies of different sizes resided within a spacious vacuole surrounded by a membrane (Fig. 6b-c). Consistent with the galectin-3 recruitment data, individual diplococci can be observed in TEM micrographs in the cytosol (Fig. 6c). Collectively, these data demonstrate that intracellular gonococcal colonies occupy a membrane-bound organelle in macrophages.

Unlike phagosomes containing diplococci and heat-killed gonococci, internalized gonococcal colonies did not accumulate early (EEA1) and late (Rab7 and LAMP2) endosomal markers (Fig 6d-e). Instead, an extensive actin filaments network surrounded the majority of intracellular gonococcal colonies within U937 MFs and primary hMDMs (Fig. 6g, Fig. S2c, and Video S1). The plasma membrane marker dsRed-mem also accumulated and co-localized with the actin network surrounding the intracellular gonococcal colonies (Fig. 6f). Collectively, these data indicate that the unique membrane-bound organelle supporting intracellular gonococcal replication is likely derived from the plasma membrane through an actin polymerization-driven process in a way that fusion with the endocytic compartment is restricted. The lack of endosomal markers on the *N*.*g* colony-containing vacuoles would suggest that colony internalization is unlikely mediated by phagocytosis, because cargo-carrying phagosomes rapidly acquire endosomal organelle identity (80).

### Gonococcal colonies invade human macrophages via a mechanism mediated by formin-dependent actin polymerization

The slow internalization kinetics as well as the absence of endosomal markers on the *N*.*g* colony-containing vacuoles are inconsistent with phagocytosis-dependent uptake and endosomal transport (65, 66). Thus, we targeted different host factors pharmacologically to identify the host determinants of *N*.*g* colony internalization. To this end, colony internalization was measured using inside/out microscopy in U937 MFs at 8hpi (Fig. 7). In parallel, we cultured U937 MFs with live non-pathogenic *Legionella pneumophila ΔdotA* (*L*.*p* Δ*dotA*) bacteria, which are taken up by non-specific phagocytosis and transported to lysosomes within minutes (71). As expected, internalization of both *L*.*p* Δ*dotA* and *N*.*g* colonies required actin polymerization (Fig. 7a, Actin^INH^). Depolymerization of microtubules with nocodazole did not affect *N*.*g* colony uptake but partially reduced phagocytosis of *L*.*p* Δ*dotA* (Fig. 7a, Tubulin^INH^). Similarly, *N*.*g* colonies were internalized efficiently when the Rho family of small GTPases were inhibited, whereas *L*.*p* Δ*dotA* phagocytosis was reduced (Fig. 7b). *L*.*p* Δ*dotA* phagocytosis required functional Cdc42 GTPase and the RhoA GTPase effector kinases ROCK1/2 and LIMK1/2, which trigger actin polymerization to shape the phagocytic cup and mediate cargo internalization (74). Phagosome closure and scission processes regulated by PI3K (69) and dynamin (70) were also required for *L*.*p* Δ*dotA* uptake (Fig. 7c). However, *N*.*g* colonies were internalized efficiently by U937 MFs even when Rho GTPases, PI3K and dynamin were inhibited (Fig. 7b-c), demonstrating that *N*.*g* colonies enter phagocytes by an actin-dependent mechanism distinct from phagocytosis.

**Figure 7.**
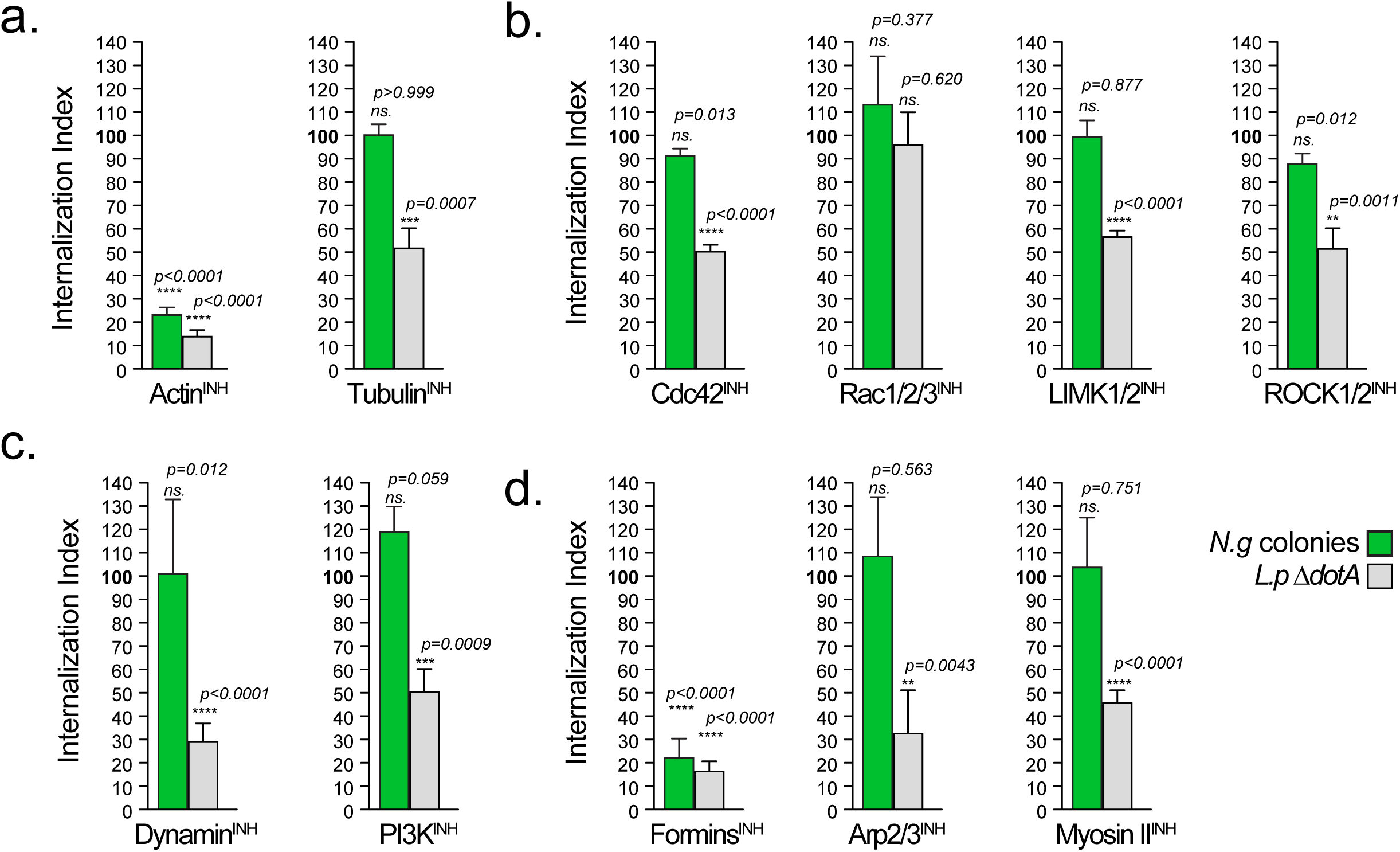
Gonococci enter macrophages despite a block in phagocytosis. Gonococci internalization by U937 macrophages requires formin-mediated actin polymerization but not phagocytosis. **(a-d)** Quantitative analysis of bacteria uptake by U937 MFs using inside/out microscopy in the presence/absence of different inhibitors added 30 min prior to infection. Gonococcal infections were stopped at 8hpi and the percentage of intracellular colonies in each condition was calculated (green bars). *L. pneumophila* Δ*dotA* infections were stopped at 2hpi and the percentage of internalized bacteria was determined (gray bars). Internalization index for each condition was calculated by dividing the percentage of internalized objects from the treatment condition by the percentage of internalized objects from the respective vehicle control condition, which was then multiplied by 100. Inhibitors targeting: **(a)** actin polymerization (cytochalasin D, 5µM) and microtubules polymerization (nocodazole, 3µM); **(b)** the small GTPases Cdc42 (ML-141, 5µM), Rac1/2/3 (EHT 1864, 5µM) and the downstream effectors of Rho GTPase-LIMK1/2 kinases (BMS-5, 1µM) and ROCK1/2 kinases (GSK269962, 1µM); **(c)** the phagosome regulators Dynamin (dynasore, 30µM) and Phosphoinositide-3 kinase (LY294002, 10µM); **(d)** actin polymerization regulators formins (SMIFH2, 25µM), Arp2/3 complex (CK869, 10µM), Myosin II ATPases ((-) blebbistatin, 5µM). Means from three biological replicates ± SD. At least 100 objects for each condition were scored. The statistical significance of the differences between the internalization index from inhibitor-treated cells and the vehicle-treated cells for each condition were calculated using the unpaired T-test.

Three main actin nucleator machineries regulate membrane morphogenesis – (1) the formin family members extend linear nuclear filaments (68); (2) the Arp2/3 complex nucleates branched actin filaments (67); (3) Myosin II ATPases which regulate the contractile actomyosin bundles (75). The formins^INH^ (68) completely blocked *N*.*g* colony internalization unlike the Arp2/3^INH^ and MyosinII^INH^ (Fig. 7d). To confirm that the Arp2/3 complex is dispensable for *N*.*g* colony uptake using an alternative approach, we utilized CRISPR/Cas9 genomic editing to generate U937 MFs in which the *ACTR2* gene encoding Arp2 is inactivated (Fig. 8c-d). Despite a general defect in phagocytosis caused by loss of Arp2, *ACTR2* KO U937 MFs internalized gonococcal colonies as efficiently as the parental cell line (Fig. 8d). Thus, gonococcal colonies elicit actin-dependent plasma membrane invaginations to invade macrophages by directly or indirectly manipulating the production of linear actin filaments mediated by one or more members of the formins family.

**Figure 8.**
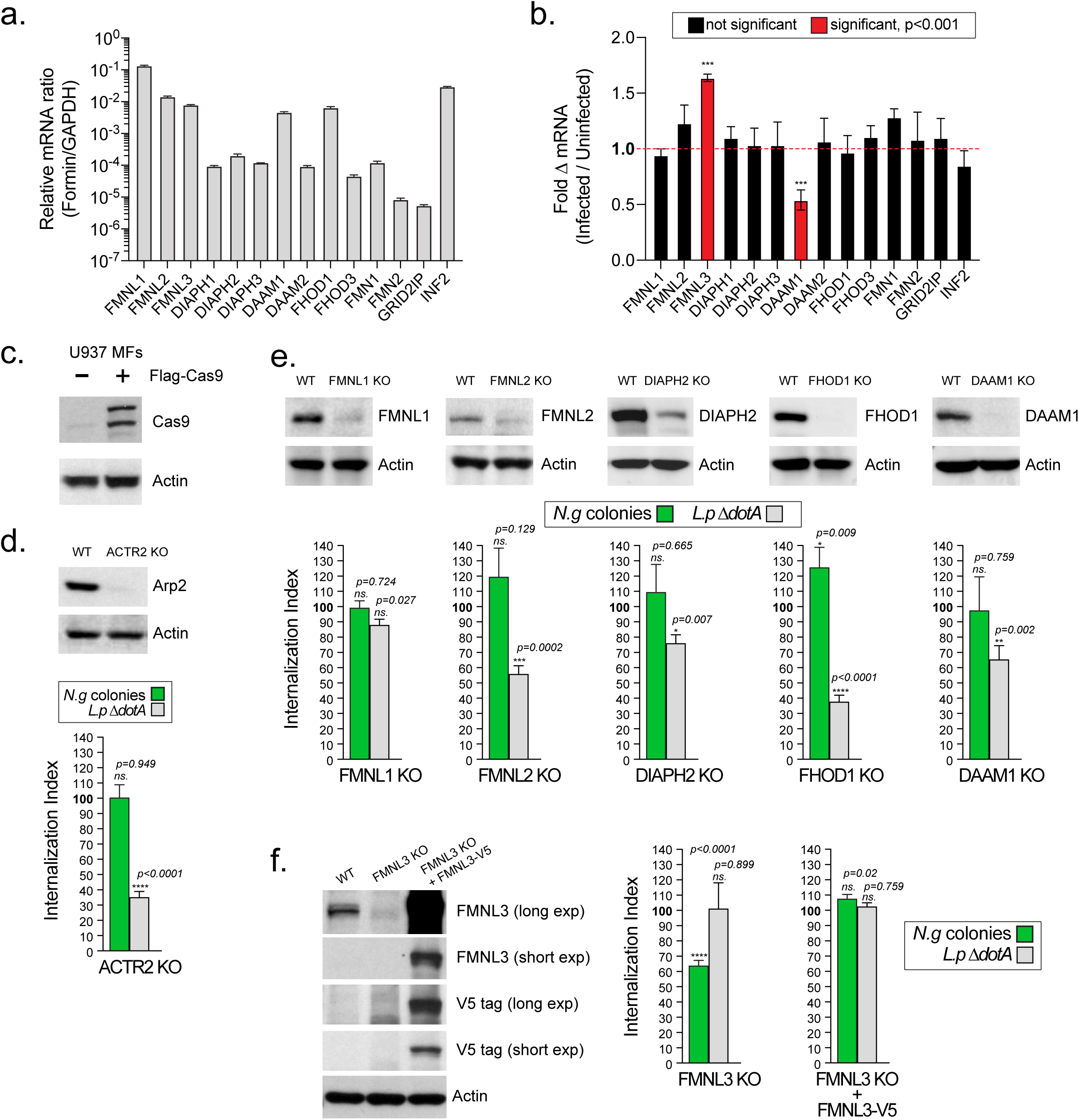
FMNL3 mediates invasion of gonococcal colonies in macrophages. **(a-b)** qPCR mRNA expression analyses for the members of the formin family in U937 MFs are shown. mRNA abundance relative to the amount of *GAPDH* mRNA is shown in **(a)**. Fold change in mRNAs abundance upon *N*.*g* FA1090 infection of U937 MFs at 4 hpi is shown in **(b). (a-b)** Means of three biological replicates ± SD are shown, *** p < 0.001 unpaired T-test. **(c)** Immunoblot analysis of the U937 cell line stably expressing Cas9 used for the CRISPR mediated genome editing. **(d)** Immunoblot shows the loss of Arp2 in the ACTR2 KO U937 MFs. Graph shows quantitative analysis of *N*.*g* colony and *Lp ΔdotA* internalization by the ACTR2 KO U937 MFs using inside/out microscopy. **(d-f)** The Internalization index for each condition was calculated by dividing the percentage of internalized objects from infections of the knock out U937 MFs cell line by the percentage of internalized objects from infections of the parental U937 MFs, which was then multiplied by 100. Values > 100% indicate increased object internalization by the KO cell line compared to WT cells, whereas values < 100% indicate decreased object internalization by the KO cell line. At least 100 objects for each condition were scored. Means from three biological replicates ± SD. The statistical significance of the differences between the internalization index from the KO cells and the parental cells for each condition were calculated using the unpaired T-test. **(e-f)** Immunoblot validation for loss of protein expression in U937 MFs differentiated from several KO cell lines in which different formin genes were inactivated. Complementation of the FMNL3 KO U937 MFs with V5-tagged FMNL3 is shown in **(f)**.

### FMNL3 localizes at sites of gonococcal invasion and mediates N.g colony internalization in human macrophages

The formin family of actin nucleators consist of fifteen members (76) – all family members except FHDC1 were expressed in U937 MFs (Fig. 8a). The mRNAs for FMNL1, FMNL2, FMNL3, DAAM1, FHOD1 and INF2 were the most abundantly expressed. Gonococcal infection did not alter the mRNA abundance of the formins family members with the exception of FMNL3 and DAAM1 (Fig. 8b). At 4 hpi the FMNL3 mRNA increased by ∼50% whereas DAAM1 mRNA abundance decreased by ∼50% (Fig. 8b). To determine which formin mediates gonococcal invasion we utilized a genetic approach to knock out individual members in U937 cells using CRISPR/Cas9 genome editing. The following U937 cell lines were successfully generated and validated via immunoblot analysis: FMNL1 KO, FMNL2 KO, FMNL3 KO, DAAM1 KO, FHOD1 KO, DIAPH2 KO (Fig. 8e-f). Next, the capacity of the individual U937 MFs knock out cell lines to phagocytose *L.p* Δ*dotA* and to internalize gonococcal colonies were compared to the parental cell line. Loss of FHOD1, FMNL2, DAAM1 and DIAPH2 significantly reduced *L.p* Δ*dotA* phagocytosis by 65%, 45%, 35% and 20% respectively but did not affect gonococcal colony internalization (Fig. 8e). Loss of FMNL3 significantly reduced gonococcal invasion by 40% despite normal *L.p* Δ*dotA* phagocytosis (Fig. 8f). Expression of C-terminally V5 tagged FMNL3 allele in the *FMNL3* KO U937 MFs fully complemented the defect in gonococcal colony internalization (Fig. 8f). These results demonstrate that gonococcal colony invasion is a process that requires FMNL3 and is mechanistically distinct from phagocytosis.

Because FMNL3 is required for gonococcal entry, presumably by mediating actin polymerization at the sites of invasion, we investigated whether FMNL3 localizes there. Antibodies against FMNL3 did not work well for immunofluorescence microscopy thus we ectopically expressed FMNL3-V5 in U937 MFs. Indeed, FMNL3 was preferentially recruited and specifically co-localized with the actin filament network at the sites of gonococcal invasion (Fig. 9a-b).

**Figure 9.**
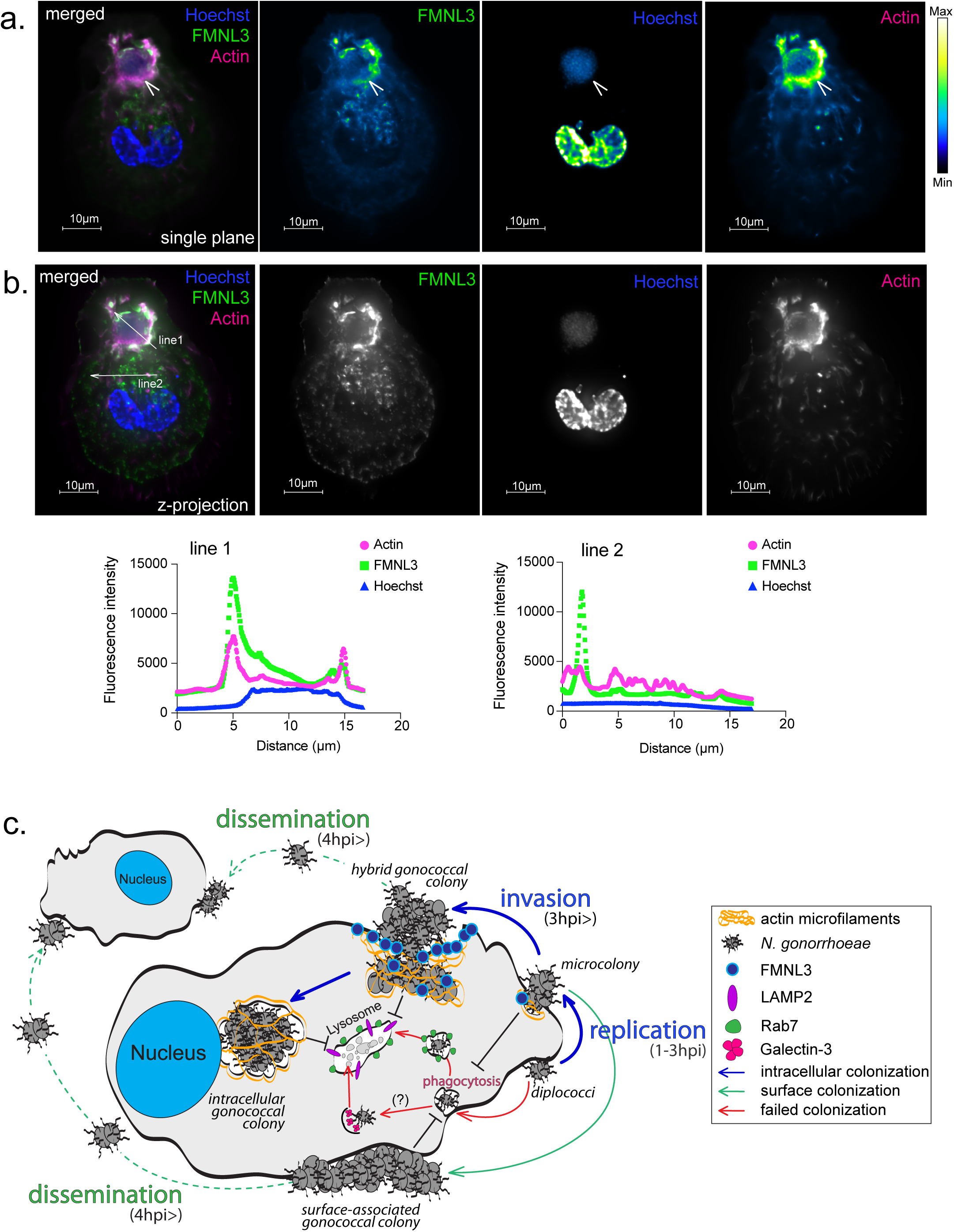
FMNL3 co-localizes with polymerized actin at the sites of *N*.*g* colony plasma membrane invasion. **(a-b)** Representative micrograph showing a *N*.*g* FA1090 colony invading a U937 MF expressing FMNL3-V5. Single plate is shown in **(a)**, z-projection is shown in **(b)**. Individual channels pseudo-colored with the Kindlmann color map that visualize fluorescent signal intensity are shown in **(a)**. Fluorescent signal profiles of **line 1** drawn along the invading colony and **line 2** drawn across another area of the cell are shown in **(b). (c)** Model of macrophage colonization and invasion by gonococcal colonies.

### CEACAM1 is dispensable for gonococcal colony invasion

Opa proteins are a phase variable multi-functional adhesins expressed by *Neisseria* species and can bind several host surface receptors (42, 47, 48, 50) to promote cell colonization as well as lipooligosaccharides (LOS) at the *N*.*g* surface to facilitate tight association among the members of the bacterial colony (77). In neutrophils infections, Opa interaction with the human CEACAM family of cell-surface receptors promotes gonococci uptake (CEACAM1, CEACAM3), trafficking to a degradative compartment (CEACAM3) and inflammatory cytokine secretion (CEACAM3) (15, 53, 54). We could not detect expression of CEACAM3 and CEACAM5 in U937 MFs by qPCR; however at least two CEACAM1 transcript variants as well as CEACAM6 were detected (Fig. S7a). Because CEACAM1 is a well-established receptor for *N*.*g* entry expressed by epithelial cells and neutrophils, we investigated if CEACAM1 acts upstream of FMNL3 to mediate *N*.*g* colony invasion. To this end, CEACAM1 KO U937 MFs were generated via genome editing (Fig. S7b). Loss of CEACAM1 did not reduce *N*.*g* colony internalization by U937 MFs indicating that CAECAM1 is dispensable for invasion and intracellular colonization of human macrophages by *N*.*g*.

## Discussion

Tissue-resident macrophages in the genitourinary tract encounter gonococci that invade the submucosa (3). Here, we show that *N*.*g* has evolved to colonize human macrophages by establishing two distinct cellular niches that independently support bacterial replication. We provide evidence for a novel colonization paradigm where invasion the host cell is carried out by an entire surface-associated bacterial colony rather than an individual bacterium. The branched multi-stage colonization model we propose in Fig. 9c defines the key stages of the macrophage-gonococci association and identifies several regulatory factors.

During the first stage (within 2hrs of contact) successful colonizers adhered to the macrophage surface, evaded phagocytosis and divided to produce microcolonies. Gonococci resisted phagocytosis by interfering specifically with their own uptake while other opsonized and non-opsonized cargo were phagocytosed normally by colonized macrophages. Gonococci that failed to block phagocytosis did not give rise to intracellular colonies but instead were transported to lysosomes and destroyed. Indeed, it has been shown that treatment with lysosome inhibitors at this early stage of the infection increased survival of gonococci within lysosomes in human macrophages (10). Thus, evasion of phagocytosis by surface gonococci in the initial stage is one critical step in macrophage colonization.

After surface microcolonies are established (2-6 hpi), the colonization process branches in the second stage where most of microcolonies begin to invade the host cell and the rest remain surface-associated. In a 24hr infection, the majority of invading colonies became intracellular and some contained hundreds of bacteria. The appearance of intracellular colonies that reside within membrane-bound organelles marks the third stage of colonization. We demonstrate that intracellular colonies originated from surface microcolonies that invade the host cell via an atypical actin-dependent mechanism rather than phagocytosis of individual diplococci. In agreement, the functions of several key phagocytosis regulators were dispensable for gonococcal invasion. Also, Arp2/3-nucleated branched microfilaments that mediate phagosome cup formation (80) were dispensable for invasion and the host membranes surrounding the invading and intracellular gonococcal colonies excluded early (EEA1) and late endosomal (Rab7) markers demonstrating that the uptake and trafficking of gonococcal colonies is distinct from the general phagocytic cargo.

Gonococci invasion required FMNL3 – a member of the formins family of actin nucleators, which assemble linear actin microfilaments that generate filopodia protrusions at the plasma membrane (81). Loss of FMNL2/3 also abolishes the protrusion forces exerted by lamellipodia without affecting the Arp2/3 complex (90). In U2OS cells FMNL3 suppression ablates filopodia formation. To our knowledge this is the first example of a bacterial pathogen entry mechanism solely based on linear actin microfilaments and does not require a functional Arp2/3 complex. Entry mechanisms by other intracellular bacteria require both linear and branched actin microfilaments (similar to lamellipodia formation) (82, 83). The uptake of *Borrelia burgdorferi* by primary human macrophages occurs through coiling phagocytosis and requires FMNL1, DIAPH1 and the Arp2/3 complex (91). Loss of FMNL3 reduced gonococcal invasion by 40%, whereas SMIFH2 blocked internalization by 80% indicating that additional family members likely also regulate macrophage entry.

Continuous bacterial replication combined with the slow invasion kinetics produces a unique topology where the gonococcal colony stretches from the surface of the macrophage and into a plasma membrane invagination that cradles a fraction of the colony. In this setting, surface gonococci acted as a plug to seal the opening of the membrane invagination thus producing a quasi-intracellular membranous compartment that is enveloped in polymerized actin microfilaments. The sharp discrete delineation observed between the protected (intracellular) and the unprotected (surface exposed) fractions of the invading bacterial colonies in our inside/out immunofluorescence studies demonstrate that gonococci and the plasma membrane are tightly opposed each other at the transition zone, which rendered gonococci within the membrane invagination protected from antibodies in the extracellular milieu. Indeed, actin microfilaments were found to bracket bacteria at the neck of the plasma membrane invagination occupied by hybrid colonies. Although our data is consistent with the notion that hybrid colonies represent a transient state during invasion, we cannot exclude the possibility that sometimes the invasion process might not be brought to completion.

We envision at least two advantages for *N*.*g* as a result of the slow invasion kinetics that produces large colonies with hybrid topology. The first is accelerated bacterial dissemination. Unlike other vacuolar intracellular pathogens that rely on cycles of invasion/replication/egress/dissemination, surface gonococci in hybrid colonies can depart to colonize neighboring bystander cells before colony invasion is completed. Indeed, we found evidence for dissemination as early as 6 hpi in macrophage infections. The second advantage is increased immune evasion plasticity. A protective niche straddling the extracellular/intracellular space allows surface-associated gonococci to escape cell autonomous defense responses targeting intracellular pathogens (such as lysosomal degradation, autophagy, and host cell death responses) (84-86) and at the same time safeguards a fraction of the colony from humoral defense factors (such as complement, circulating antibodies and anti-microbial peptides) (87). Therefore, the gonococci adaptation to multiple topologically distinct niches could be an elegant evolutionary solution to simultaneously counteracts multiple inherently distinct host defense responses albeit at the cost of protecting only a fraction of the colony members.

The mechanism that blocks invasion and allows a fraction of gonococcal colonies (∼20%) to remain at the cell surface and expand is unclear. One reason could be loss of expression of a gonococcal invasin. *Neisseria* species extensively utilize phase and antigen variation to remodel the bacterial surface composition as an immune evasion strategy to increase antigenic diversity (20-23). Alternatively, gonococci-driven host cell reprogramming might be required to remove an invasion block imposed by the colonized macrophage. Future experimental work is needed to distinguish between the two invasion models.

Infrequently, we detected individual diplococci in the cytosol of infected macrophages that were targeted to autophagosomes as indicated by the accumulation of galectin-3 and LC-3. Gonococci have been reported to localize in the cytosol of epithelial cells (31, 32) and macrophages (10); thus, it has been speculated that bacterial replication takes place in the macrophage cytosol. However, our studies demonstrated that in macrophage infections gonococci replicate intracellularly within a membrane-bound organelle. Rupture of bacterium-containing phagosome can deposit vacuolar pathogens in the host cytosol, which are subsequently cleared by autophagy (84, 85). It is possible that cytosolic escape is a byproduct of a defect in entry, which would explain why single bacteria, but not gonococcal colonies localized to the cytosol. However, we cannot exclude the possibility that gonococci encode the capacity to replicate in the cytosol of eukaryotic cells. Potentially, macrophages might be more efficient at eliminating cytosolic bacteria and thus prevent gonococci from replicating in the cytosol. Collectively, our data highlights that gonococci likely have evolved to establish multiple distinct cellular niches simultaneously likely to overcome cell intrinsic defense responses.

Several aspects of macrophage colonization are distinct from how the gonococci interacts with other host cell types encountered during infection. Both gonococci and meningococci form epicellular colonies on epithelial cells, in which the base of the colony is partially embedded in actin-rich plasma membrane ruffles known as actin plaques (56). Actin plaques accumulate Ezrin - an adaptor protein that cross-links cell surface receptors with the cortical actin network (57-60). Gonococcal surface colonies did not produce actin plaques on macrophages (Fig S2c) and Ezrin was absent from the actin cages that surround the intracellular colonies (Fig S6e). Thus, the actin subversion mechanisms utilized by gonococci for macrophages and epithelial cells colonization are distinct. Also, gonococci were found to be both intracellular and surface-associated in neutrophil infections (31-33, 53). The uptake of gonococci by neutrophils is strictly species specific and is mediated by the engagement of the Opa adhesins with the human CEACAM1/CEACAM3 surface receptors (78, 79). However, our data demonstrates U937 MFs did not express CEACAM3 and that CEACAM1 is dispensable for macrophage entry.

The four different gonococcal strains we tested colonized U937 MFs and primary human MDMs similarly and produced intracellular as well as extracellular colonies. The actin accumulation on the gonococci-containing vacuoles was indistinguishable in both cell types. Thus, U937 macrophages in many aspects of gonococcal infections recapitulate human primary monocyte-derived macrophages.

Although the initial phases of *N*.*g* infections and its interactions with mucosal epithelium and neutrophils have been extensively studied, the cellular niche in the submucosa that supports *N*.*g* growth *in vivo* is still poorly understood. Considering the short neutrophil lifespan (< 24hrs), which gonococci can extend by a few hours (88, 89), our results argue that macrophages could represent an underappreciated cell type that can support rapid gonococcal expansion in the submucosa for longer time periods (> 48hrs).

## Material and Methods

### Bacterial strains and culture conditions

*Neisseria gonorrhoeae* isolates FA1090, F62, MS11 and F19 were used in this study. All *Neisseria* strains were kind a gift from Dr. Hank Seifert (Northwestern University). Bacteria were cultured on gonococcal medium base (GCB) agar (Criterion) with Kellog’s supplements (92) at 37°C and 5% CO_2_ for approximately 16 hrs. For infections, 4 to 6 piliated Opa+ colonies were pooled, streaked on GCB agar (1×1cm patch) and grown for 16hrs. Patches were collected, resuspended in PBSG (PBS supplemented with 7.5mM Glucose, 0.9 mM CaCl_2_, and 0.7 mM MgCl_2_) and the number of bacteria were determined by OD_550_ measurements. Inoculums for infections were prepared in PBSG and the number of bacteria were confirmed by plating dilutions from the inoculum on GCB agar. For experiments with heat killed bacteria, the inoculum was split, and one set was heat-killed at 70°C for 30min. Loss viability was confirmed by plating the bacteria on GCB agar.

*Legionella pneumophila* serogroup 1 strain LP01 with a clean deletion of the *dotA* gene was used in this study (71). The *Legionella* strain was grown on charcoal yeast extract (CYE) plates [1% yeast extract, 1%N-(2-acetamido)-2-aminoethanesulphonic acid (ACES; pH 6.9), 3.3 mM l-cysteine, 0.33 mM Fe(NO3)3, 1.5% bacto-agar, 0.2% activated charcoal] (93). For the macrophage phagocytosis studies, *Legionella* were harvested from CYE plates after growth for 2 days at 37°C, resuspended in H_2_O and bacterial number was determined by OD_550_ measurement.

### Cells and culture conditions

The human monocytic cell line U937 (ATCC CRL-1593.2) was obtained from ATCC and primary human peripheral monocytic cells (PBMCs) from two different donors were purchased from Precision for Medicine. U937 cells were cultured in RPMI 1640 media (VWR) supplemented with 10% Fetal Bovine Serum (FBS) at 37°C and 5% CO_2_. For differentiation into mature macrophages, U937 monocytes were seeded at the desired density, cultured with 10 ng/ml phorbol 12-myristate 13-acetate (PMA) for 24hrs followed by 48hrs incubation with RPMI media (+10% FBS) in the absence of PMA. PBMCs were differentiated into mature macrophages with human recombinant macrophage colony stimulating factor (BioVision) in RPMI 1640 media supplemented with 20% human AB serum (Corning) for 7 days.

### Antibodies and Inhibitors

The purified chicken IgY anti-*Neisseria gonorrhoeae* antibody was raised against formalin-killed FA1090 gonococci by Cocalico Biologicals and it cross-reacts with the MS-11, FA19 and F62 strains. The rabbit anti-*N*.*g* polyclonal antibody (cat# 20-NR08) was purchased from Fitzgerald Industries. Rabbit monoclonal antibodies against the following antigens were purchased from Cell Signaling - Rab5 (C8B1), Rab7 (D95F2), EEA1 (C45B10), GM130 (D6B1), Ezrin (#3145), LC3A/B (D3U4C), V5-Tag (E9H80), CEACAM1 (D3R80) and ACTR2 (#3128S). Mouse antibodies against the following antigens were purchased from Santa Cruz – Actin (sc8432), FMNL1 (sc390466), FMNL2 (sc390298), FHOD1 (sc365473), DAAM1 (sc100942). Mouse anti-Cas9 (7A9-3A3) was purchased from Cell Signaling. Anti-FMNL3 (ab57963) and anti-Galectin 3 (ab2785) mouse monoclonal antibodies were purchased from Abcam. Anti-human LAMP-2 (H4B4) was purchased from BioLegend. Anti-DIAPH2 (A300-079A-T) was purchased from Bethyl Labs. Highly cross-absorbed Alexa fluorophore conjugated secondary antibodies were purchased from Life Technologies.

Inhibitors used in this study were purchased from: (1) Cayman Chemicals - cytochalasin D (5µM), CK869 (10µM), Dynasore (30µM), GSK269962 (1µM), Nocodazole (3µM), ML-141 (5µM), EHT 1864 (5µM), BMS-5 (1µM) and (-) blebbistatin (5µM); (2) Cell Signaling - LY294002 (10µM); (3) Abcam - SMIFH2 (25µM).

### Bacterial Survival assays

Macrophages differentiated in 24-well plates at 2×10^5^ cells/well were infected with an inoculum of gonococci grown as a patch from 4-6 piliated colonies on GCB agar for 16hrs. The patch was collected in pre-warmed PBSG medium, the bacterial number was determined by OD_550_ densitometry and the MOI of the inoculum was at 1 bacterium per 10 macrophages. Bacteria were added to macrophages-containing wells in PBSG or empty wells on the same plate. The plate was centrifuged for 5min at 1000 rpm to bring the bacteria in contact with the macrophages. The T_0_ CFU counts were determined by plating a dilution of the inoculum on GCB agar. As indicated in each experiment, at different times post infection the number of the gonococci in the supernatant and in cells lysed by directly lysing the eukaryotic cells with 0.05% Tween-20 (5 min; 37C) and serial dilutions were plated on GCB agar to determine CFU counts. Three technical replicates were done for each condition in each experiment.

### Contact-dependent bacterial replication assay

Macrophages differentiated in 6-well plates at 1×10^6^ cells/well were infected with an inoculum of gonococci grown on GCB agar for 16hrs as a patch started from 4 to 6 piliated colonies. The patch was collected in pre-warmed PBSG medium, the bacterial number was determined by OD_550_ densitometry and the MOI of the inoculum was at 2 bacteria per macrophage. The inoculum was prepared with PBSG and the infection was carried out in PBSG. Transwell inserts with a 0.4µm pore size bottom membranes (Corning) were placed in each well. The inoculum was added either within the transwell insert to measure contact independent bacterial replication or in the well to measure contact dependent replication. To establish a baseline of gonococcal replication in PBSG the inoculum was also added to wells that did not contain macrophages. The plate was centrifuged for 5min at 1000 rpm to bring the bacteria in contact with the macrophages. The T_0_ CFU counts were determined by plating a dilution of the inoculum on GCB agar. As indicated in each experiment, at different times post infection the number of the gonococci in the supernatant and in cells lysed by directly lysing the eukaryotic cells with 0.05% Tween-20 (5 min; 37C) and serial dilutions were plated on GCB agar to determine CFU counts. To confirm the transwell membrane integrity in the contact independent growth conditions, macrophages were lysed with detergent, serial dilutions were plated on GCB agar, and as expected viable CFU were not recovered. Three technical replicates were performed for each condition in each experiment.

### Immunofluorescence microscopy of infected macrophages

For infections, U937 monocytes were PMA-differentiated directly on cover slips in 24-well plates whereas hMDMs were seed on cover slips in media (RPMI + 10% FBS) for 20hrs prior to infection. The seeding density for all cell types was kept at 2×10^5^ cells/well. The MOIs used for each experiment and the duration of infection are indicated in the Figure legends. All infections were carried out in PBSG. We did not observe increase in macrophage cell death even after prolonged (> 24hrs) culture with PBSG. After the inoculum addition, cell culture plates were spun down at 1,000 rpms for 5min to facilitate bacteria-macrophage contact.

For immunofluorescence microscopy analysis, coverslips were washed with PBS three times, fixed with 2% paraformaldehyde (PFA) for 60 min at ambient temperature, permeabilized with 0.1% Triton X-100 for 20min, and blocked with 2% goat serum in PBS for 60 min. Next, primary antibodies were added in PBS containing 0.2 % goat serum either overnight at 4°C or for 90min at room temperature. The antibodies dilutions were 1:500 for the α-*N*.*g* antibodies and 1:200 for all other primary antibodies. Secondary antibodies were used at 1:500 dilution and Hoechst 33342 at 1:2000 for 90 min at room temperature. The high affinity actin probe Alexa Fluor-568 Phalloidin (ThermoFisher) was used at 1:2000 dilution and was added with the secondary antibodies. Coverslips were mounted with ProLong Gold antifade reagent (ThermoFisher) onto slides and examined by fluorescence microscopy.

The different bacterial sub-populations in the inside/out staining methodology were detected by differential immunolabeling before and after detergent permeabilization of host cell the plasma membrane. Briefly, extracellular gonococci were immunolabeled with rabbit α-*N*.*g* antibody (Fitzgerald Ind.) in 2% goat serum for 90 min after the infected cells were washed with PBS three times, fixed with 2% PFA for 60 min at ambient temperature. The antibody was immobilized on the samples with 2% PFA (60 min incubation). Next, all bacteria were labeled with chicken α-*N*.*g* IgY antibody (Cocalico Biologicals) for 90 min after the samples were permeabilized with 0.1% Triton X-100 for 20min and blocked with 2% goat serum in PBS for 60 min. In each micrograph, antibody inaccessible gonococci were single positive whereas the surface associated bacteria were double positive.

Microscopy analyses of infected cells. Images were acquired with inverted wide-field microscope (Nikon Eclipse Ti) controlled by NES Elements v4.3 imaging software (Nikon) using a 60X/1.40 oil objective (Nikon Plan Apo λ), LED illumination (Lumencor) and CoolSNAP MYO CCD camera. Image acquisition and analysis was completed with NES Elements v4.3 imaging software. Only linear image corrections in brightness or contrast were completed. For all analyses, three-dimensional images of randomly selected fields were acquired and image acquisition parameters were kept constant for all cover slips from the same experiment. The Z depth acquisition was set based on the out-of-focus boundaries and the distance between individual Z-slices was kept at 0.3µm.

### Analysis of gonococcal invasion

U937 macrophages differentiated on cover slips were pre-treated with the following inhibitors for 30 min prior to infection: cytochalasin D (5µM), CK869 (10µM), SMIFH2 (25µM), GSK269962 (1µM), LY294002 (10µM), Nocodazole (3µM), ML-141 (5µM), EHT1864 (5µM), BMS-5 (1µM), Dynasore (30µM) and (-) blebbistatin (5µM).

The inhibitors remained for the duration of the infection. Macrophages were infected with *N*.*g* FA1090 (MOI = 2) for 8hrs or with *L. pneumophila ΔdotA* (MOI = 40) for 2hrs, after which the cells were extensively washed with PBS and fixed with 2% PFA. Samples were processed for inside/out immunofluorescence analysis (as detailed in the previous section) and the number of internalized bacteria were determined by microscopy analysis.

### Bacteria quantitation using 3D immunofluorescence microscopy

To calculate the number of gonococci within a colony or the number of bacteria populating each host cell in 3D micrographs of gonococci-colonized macrophages we utilized the methodology we previously developed to count bacterial cells from *Legionella*-infected macrophages (94). Briefly, 3D images (Z-slice = 0.3µm) of infected cells were analyzed by setting a binary mask using bacterial fluorescence to define individual objects. Within each object, the number of bacteria varied from individual diplococci to hundreds of bacteria (in large colonies). The formula to calculate the number of bacteria within an object volumes (***X****=0*.*4093\*****V****+1*.*134*, **R**^**2**^=0.9037, **X**-number of gonococci, **V**-object volume [voxels]) was derived from volume measurements data of individual bacteria as well as bacterial colonies (n = 142) where bacteria can be unambiguously identified and counted manually. Cortical actin was used to delineate the boundaries of individual infected macrophages.

### Macrophage phagocytosis assays

U937 macrophages differentiated on cover slips were infected with *N*.*g* FA1090 (MOI = 2) for 7hrs. Next, phagocytic cargo of live bacteria (MOI = 40) or latex particles were added for 60 min after which the cells were extensively washed with PBS and fixed with PFA. To measure FcR-mediated phagocytosis FITC-labeled rabbit IgGs conjugated to latex beads with 0.1µm mean particle size (Cayman Chemicals) were used as cargo. The *Legionella pneumophila* Δ*dotA* was used to measure phagocytosis of live bacteria because this mutant is avirulent, is taken up via phagocytosis, traffics through the endocytic compartment and is killed by lysosomal degradation (71). To determine cargo internalization, an inside/out staining methodology was utilized as follows, samples were: (1) fixed with 2% PFA (60 min), washed and blocked with 2% goat serum in PBS; (2) incubated with rabbit anti-*N*.*g* antibody (Fitzgerald Ind.) in the presence (Δ*dotA* infections) or absence (FITC-IgG latex beads treatment) of rabbit anti-*Legionella* antibody (Cocalico) for 60min; (3) fixed with 2% PFA (60 min), washed and permeabilized with 0.1% Triton X-100 for 20min; (4) incubated with purified chicken anti-*N*.*g* IgY antibody (Cocalico) in the presence (Δ*dotA* infections) or absence (FITC-IgG latex beads treatment) of purified chicken anti-*Legionella* IgY antibody (Cocalico) for 60min; (5) washed extensively and incubated with highly cross-absorbed anti-rabbit Alexa568 (ThermoFisher), anti-chicken IgY FITC (Life Technologies), and Hoescht 33342 (Molecular Probes) for 60 min. Size and morphology were used as determinants to differentiate between *Neisseria* (diplococcus, ∼1µm), *Legionella* (rod, ∼1.5µm) and the latex beads (amorphous aggregates).

### Thin-section transmission electron microscopy

Differentiated U937 macrophages were infected with *N*.*g* FA1090 for 10hrs (MOI=2) on cover slips. The cells were washed with PBS and fixed with 1.6% glutaraldehyde and 2.5% paraformaldehyde in 0.1M Cacodylate Buffer for 60 min at ambient temperature. The samples were processed and imaged at the LSU School of Veterinary Medicine Imaging Core facilities (https://www.lsu.edu/vetmed/cbs/facilities/microscopy_center/index.php) on a JEOL JEM-1011 transmission electron microscope.

### Quantitative Real-time PCR analysis

U937 macrophages seeded in 24-well plates (2×10^5^ cells/well) were infected with *N*.*g* FA1090 for 8hrs (MOI=2). In parallel, samples from uninfected cell were also collected. Total RNA isolation and first-strand cDNA synthesis was performed using TaqMan Gene Expression Cells-To-Ct Kit (ThermoFisher) as recommended by the manufacturer. Amount of the different mRNAs were determined by quantitative real-time PCR using the Universal ProbeLibrary (Roche, Life Science) and LightCycler 480 Probes Master (Roche, Life Science). The primers detecting the different CEACAMs mRNAs and their splice variants were designed by the Universal ProbeLibrary Assay Design Center (Roche) and are listed in Supplementary Table 1. Thermal cycling was carried out using a LightCycler 96 instrument (Roche Diagnostics) under the following conditions: 95°C for 5 minutes and 45 cycles at 95°C for 10s and 60°C for 25s. Gene expression was normalized to GAPDH.

### Generation of U937 stable cell lines

Formins knockout U937 cell lines were generated using lentiCas9-Blast (Addgene plasmid 52962) and lentiGuide-Puro (Addgene plasmid 52963) as previously described (95). Guide RNAs used in this study are listed in Supplementary Table 2.

FMNL3-V5 for complementation studies was purchased from Arizona State University Biodesign Institute DNASU Plasmid Repository. FMNL3-V5 was expressed in U937 FMNL3 KO using lentiviruses generated in 293E cells co-transfected with the packaging constructs psPAX2 (Addgene plasmid 12260) and pCMV-VSV-G (Addgene plasmid 8454).

U937 cell lines stably expressing red fluorescent protein membrane were generated using pDsRed-Monomer-Mem (Takara Bio) cloned into pLB vector from Addgene (Addgene plasmid 11619) as previously described (96). The corresponding lentiviruses were generated in 293E cells co-transfected with the packaging constructs psPAX2 and pCMV-VSV-G as described.

### Statistical analysis

Calculations for statistical differences were completed by Student’s T-test as indicated using Prism v9 (GraphPad Software).

## Author Contributions

AMD and SSI conceptualized the project and acquired funds for the project. RC, MC, SSI and AMD performed the experimental work and data analysis. AMD and SSI wrote the original draft, reviewed and edited the final manuscript.

## ACKNOWLEDGEMENTS

We are grateful to Dr. Hank Seifert (Northwestern University) for providing the *Neisseria* strains used in this study and to Dr. Xiaochu Wu (LSU School of Veterinary Medicine) for technical assistance with processing samples for TEM and image acquisition. This work was supported by start-up funds from LSU Health Sciences Center-Shreveport to SSI and to AMD and by U54 GM104940 from the National Institute of General Medical Sciences of the National Institutes of Health, which funds the Louisiana Clinical and Translational Science Center (LACaTS). The content is solely the responsibility of the authors and does not necessarily represent the official views of the National Institutes of Health.

