## Supplemental Material for "*Neisseria gonorrhoeae* subverts formin-dependent actin polymerization to colonize human macrophages"

#### SUPPLEMENTAL FIGURES LEGENDS

**Figure S1. Kinetics of U937 MFs infections with *N.g* FA1090.** Micrographs showing representative fields of macrophages infected with *N.g* FA1090 (MOI = 0.1) over a 22hrs infection. Gonococci were detected with polyclonal chicken anti-*Ng* IgY antibody. Hoechst labels the macrophage nucleus. The arrowhead points to a super-infected macrophage at 22 hpi which contains hundreds of gonococci.

**Figure S2. Distinct actin assembly at gonococcal colonies on hMDMs and epithelial cells. (a)** Representative micrograph showing an actin plaque underneath a surface-associated colony at 8hpi following HeLa229 cells infections with *N.g* FA1090. **(b)** Quantitative analysis of actin plaque formation underneath surface-associated colonies on Hela cells and U937 MFs infected with *N.g* FA1090 for 8 hours. Means of technical triplicates ± SD are shown, ** $p < 0.005$ unpaired T-test. **(c)** Micrograph of human MDMs infected with FA1090 at 8hpi showing representative neighboring macrophages that harbor either a hybrid (arrowhead) or a surface-associated colony (open triangle). Honeycomb pattern shaped acting cages assemble around intracellular regions of the hybrid colony but actin plaques did not form underneath the surface colony. Individual focal plains of a 2µm Z-stack are shown. The actin channel is shown individually in grayscale as well. (**d**) Micrograph (Z-stack projection) of a hybrid colony on a hMDM infected with *N.g* FA1090 at 8hpi that is also shown in 3D in the supplementary movie 1. **(a, c-d)** Colony topologies were determined by inside/out 3D microscopy.

**Figure S3. Quantitative analysis of gonococcal uptake by U937 MFs infected with *N.g* FA1090.** The localization for each bacterium was scored based on inside/out microscopy of infected cells as indicated. **(a)** Percentage of live and heat-killed gonococci internalized by macrophages at 1hpi (MOI=2). **(b)** Presence of gonococcal microcolonies (4 to 12 bacteria) at 2hpi on U937 MFs infected at MOI=2 with live or heat-killed bacteria. (**c**) Analysis of the topology of U937 MFs-associated gonococcal microcolonies is shown. **(a-c)** Means of technical triplicates ± SD ** $p < 0.005$ unpaired T-test. At least 100 object were scored for each condition.

**Figure S4. Opsonized and non-opsonized cargo uptake by gonococci colonized U937 MFs. (a-c)** Representative micrographs of U937 MFs pulsed with opsonized IgG-coated latex beads (0.1 µm mean particle size) or non-opsonized live *L.p* Δ*dotA* bacteria (MOI=40) for 60min. Z-stack projections of inside/out stained samples are shown. (**a**) IgG-beads uptake by uninfected U937 macrophages. **(b-c)** Cargo uptake by U937 macrophages colonized with gonococci for 7hrs (*N.g* FA1090). **(d)** Quantitative analysis of cargo uptake for 60min by uninfected macrophages vs. macrophages colonized with *N.g* FA1090. Data shows the percentage of uninfected macrophages with internalized cargo as percentage of all uninfected cells vs. the percentage of surface colonized macrophages with internalized cargo as percentage of all surface colonized macrophages. 3D inside out microscopy was used to determine gonococci and cargo location. Size and morphology were used as determinants to differentiate between *Neisseria* (diplococcus, ~1µm), *Legionella* (rod, ~1.5µm) and the latex beads (amorphous aggregates). Means of technical replicates ± SD are shown, at least 50 cells were analyzed for each condition.

**Figure S5. Quantitative analysis of intracellular and surface-associated gonococcal replication. (a-b)** Surface-associated replication of *N.g* FA1090 on U937 MFs pre-treated with 5µM cytochalasin D or with DMSO. **(a)** Representative micrographs of large surface associated colonies at 8hpi on U937 MFs pre-treated with 5µM cytochalasin D. **(b)** Gonococcal CFUs recovered from 8 hour infections (MOI = 0.1) in the presence/absence of 5µM cytochalasin D. **(c-d)** Quantitative analysis of gonococcal invasion **(c)** and size of gonococcal colonies **(d)** at 10 hpi in U937 MFs infections when cells were either treated with DMSO or cytochalasin D at 4 hpi. Inside/out microscopy was used to determine the colony topology and object volume measurement was used to determine the size of each macrophage-associated *N.g* colony. Each data point represents an individual colony. At least 30 objects for each condition were analyzed. Means ± SD are shown, unpaired T-test

**Figure S6. Cellular distribution of different host proteins in U937 MFs colonized by *N.g* FA1090 gonococci. (a-e)** Single focal plane representative micrographs from 3D images showing subcellular localization of (a) Rab7, (b) LAMP2, (c) Rab5, (d) LC3 and (e) Ezrin.

**Figure S7. Gonococci invasion of human macrophages is independent of CEACAM1. (a)** qPCR mRNA analysis for *ceacam1*, *ceacam3*, *ceacam5* and *ceacam6* was carried out on U937 MFs that were either infected with *N.g* FA1090 for 8hr (MOI=2) or were left uninfected. Primer sets specific for different transcripts splice variants (*tv*) were designed and used when possible. Means of biological triplicates ± SD are shown. **(b)** Immunoblot shows the loss of CEACAM1 in the CEACAM1 KO U937 MFs. **(c)** Graph shows quantitative analysis of *N.g* colony and *Lp* Δ*dotA* internalization by the CEACAM1 KO U937 MFs using inside/out microscopy. The Internalization index for each condition was calculated by dividing the percentage of internalized objects from infections of the CEACAM1 KO U937 MFs cell line by the percentage of internalized objects from infections of the parental U937 MFs, which was then multiplied by 100. Values > 100% indicate increased object internalization by the KO cell line compared to WT cells, whereas values < 100% indicate decreased object internalization by the KO cell line. At least 100 objects for each condition were scored. Means from three biological replicates ± SD. The statistical significance of the differences between the internalization index from the KO cells and the parental cells for each condition were calculated using the unpaired T-test.

**Supplemental Video 1. Representative 3D micrograph of a hybrid gonococcal colony invading a human primary monocyte-derived macrophage.** The movie is assembled from sequential single focal plane micrographs along the Z-axis which are separated by 0.3µm from each other. The video shows a macrophage infected with *N.g* FA1090 that is processed for inside/out microscopy. Purple arrow indicates the gonococcal colony. White arrows point to the surface exposed bacteria from the colony, which are false colored green.

Figure S1.

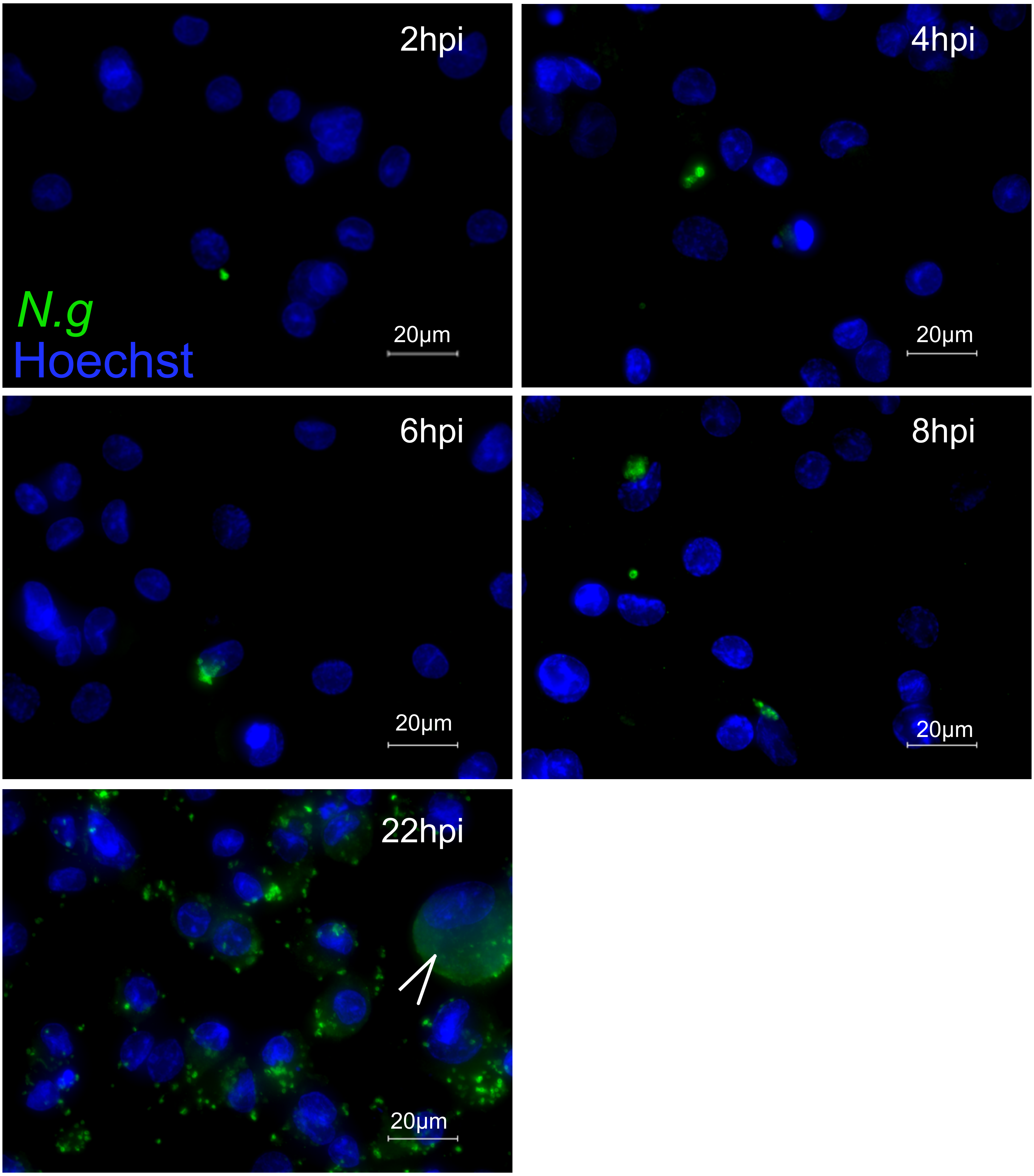

### Figure S2.

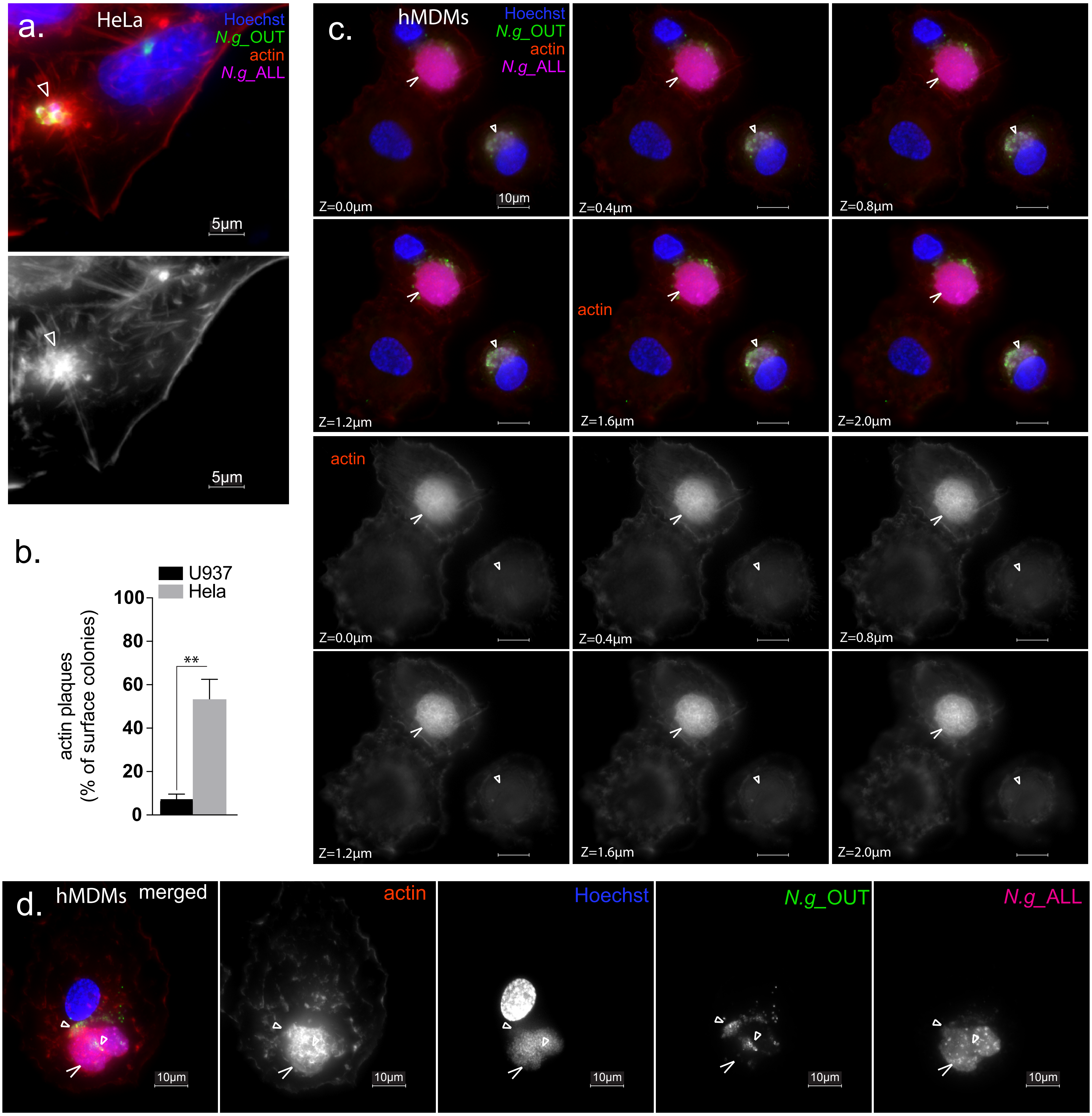
a.
HeLa
Hoechst
N.g_OUT
actin
N.g_ALL
5µm
5µm
b.
U937
Hela
actin plaques
(% of surface colonies)
100
80
60
40
20
0
**
c.
hMDMs
Hoechst
N.g_OUT
actin
N.g_ALL
10µm
Z=0.0µm
Z=0.4µm
Z=0.8µm
Z=1.2µm
Z=1.6µm
Z=2.0µm
actin
actin
Z=0.0µm
Z=0.4µm
Z=0.8µm
Z=1.2µm
Z=1.6µm
Z=2.0µm
d.
hMDMs
merged
actin
Hoechst
N.g_OUT
N.g_ALL
10µm
10µm
10µm
10µm
10µm

### Figure S3.

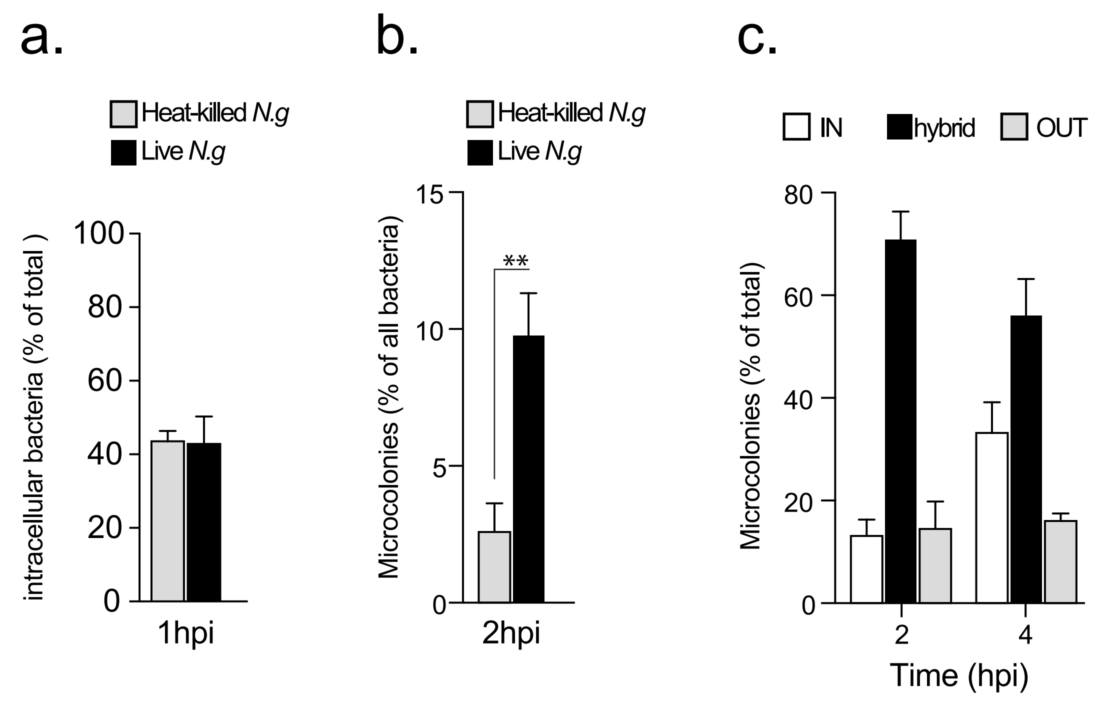
a.
Heat-killed N.g
Live N.g
intracellular bacteria (% of total )
100
80
60
40
20
0
1hpi
b.
Heat-killed N.g
Live N.g
Microcolonies (% of all bacteria)
15
10
5
0
**
2hpi
c.
IN
hybrid
OUT
Microcolonies (% of total)
80
60
40
20
0
2
4
Time (hpi)

Figure S4.

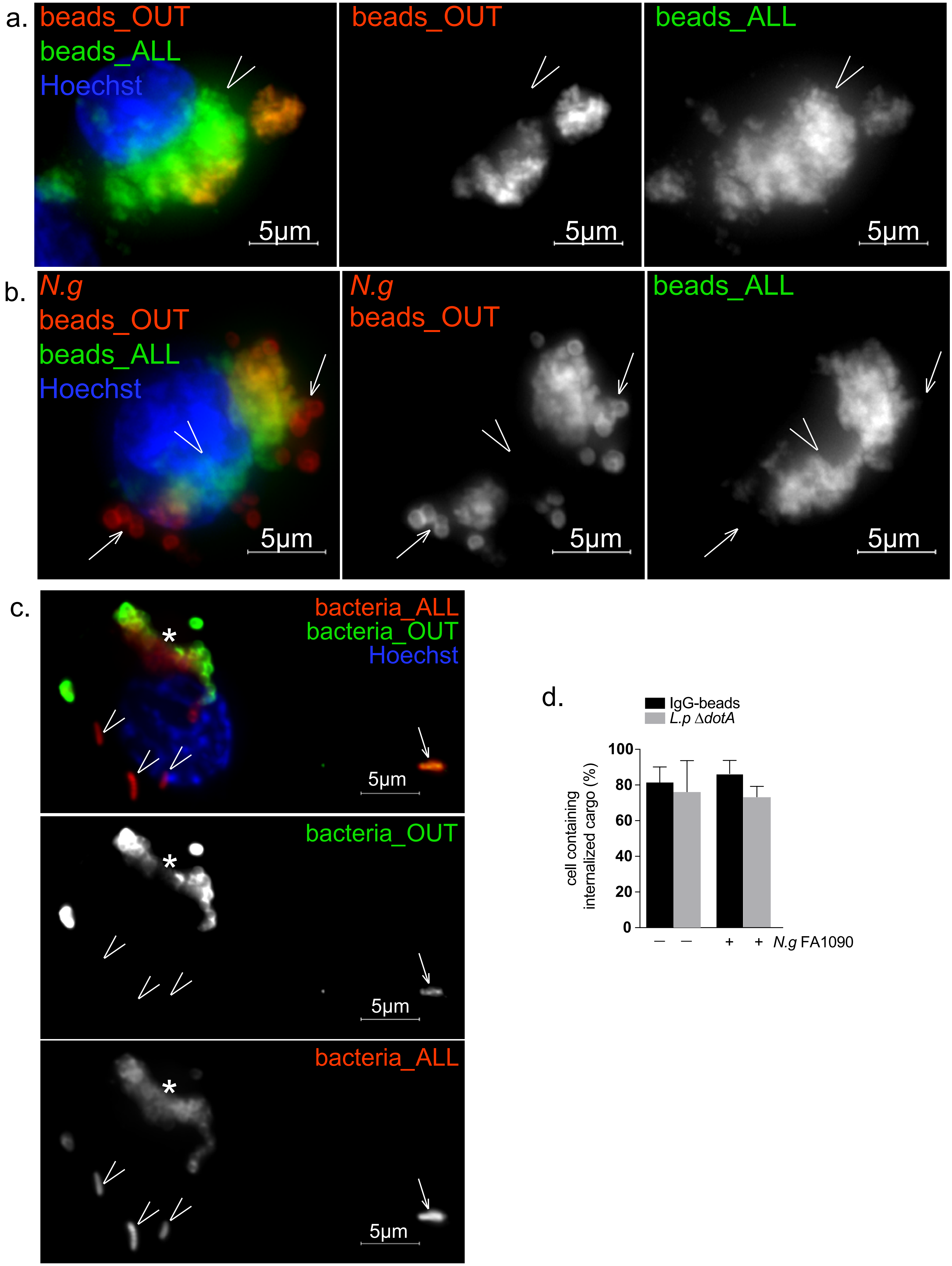
a.
beads_OUT
beads_ALL
Hoechst
5µm
beads_OUT
5µm
beads_ALL
5µm
b.
N.g
beads_OUT
beads_ALL
Hoechst
5µm
N.g
beads_OUT
5µm
beads_ALL
5µm
c.
bacteria_ALL
bacteria_OUT
Hoechst
*
5µm
bacteria_OUT
*
5µm
bacteria_ALL
*
5µm
d.
IgG-beads
L.p ΔdotA
cell containing internalized cargo (%)
0
20
40
60
80
100
–
–
+
+
N.g FA1090

Figure S5.

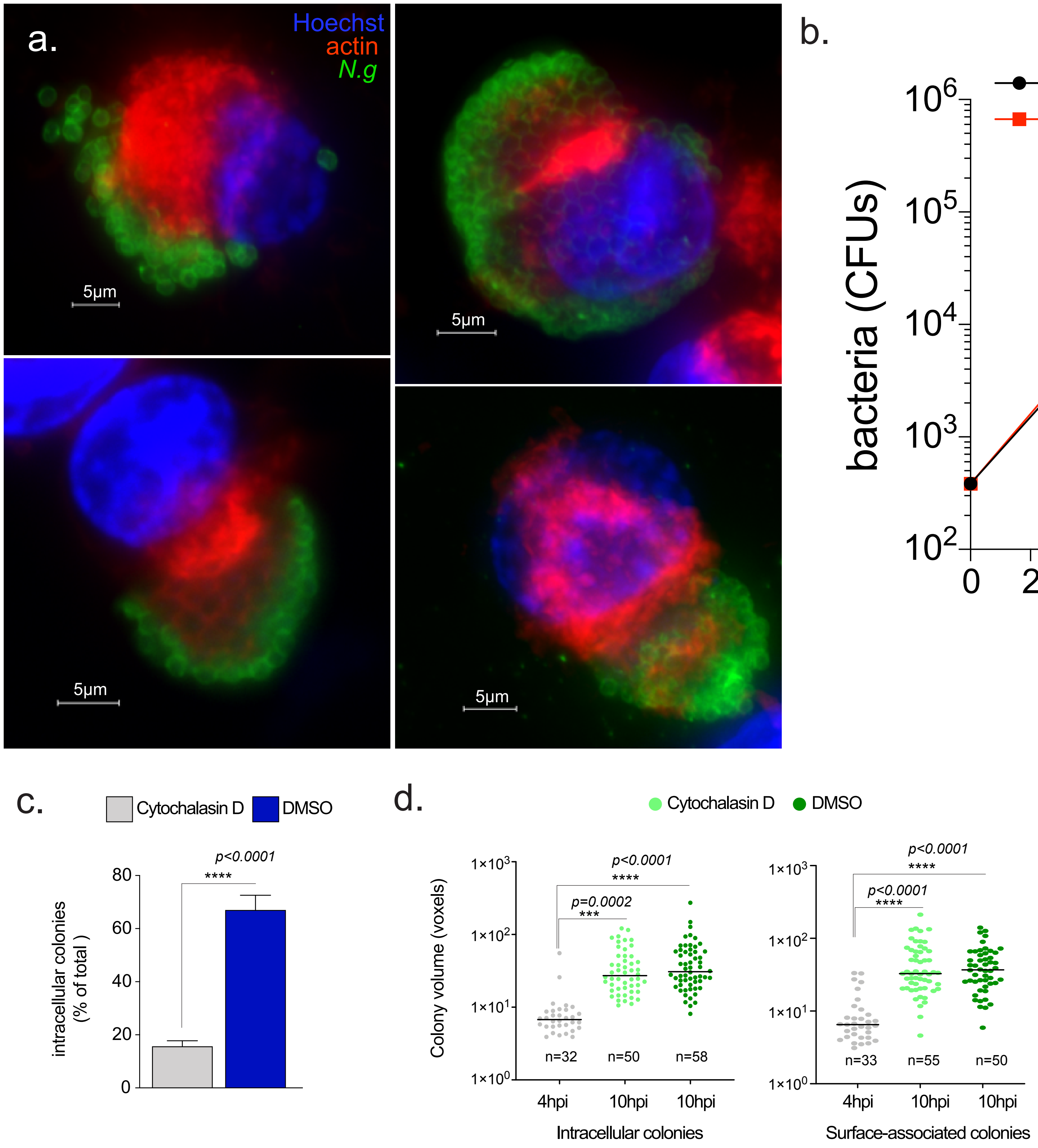
a.
Hoechst
actin
N.g
5μm
5μm
5μm
5μm
b.
DMSO
Cyto D
bacteria (CFUs)
Time (hpi)
c.
Cytochalasin D
DMSO
intracellular colonies (% of total )
p<0.0001
****
d.
Cytochalasin D
DMSO
Colony volume (voxels)
p<0.0001
****
p=0.0002
***
n=32
n=50
n=58
4hpi
10hpi
10hpi
Intracellular colonies
p<0.0001
****
p<0.0001
****
n=33
n=55
n=50
4hpi
10hpi
10hpi
Surface-associated colonies

Figure S6.

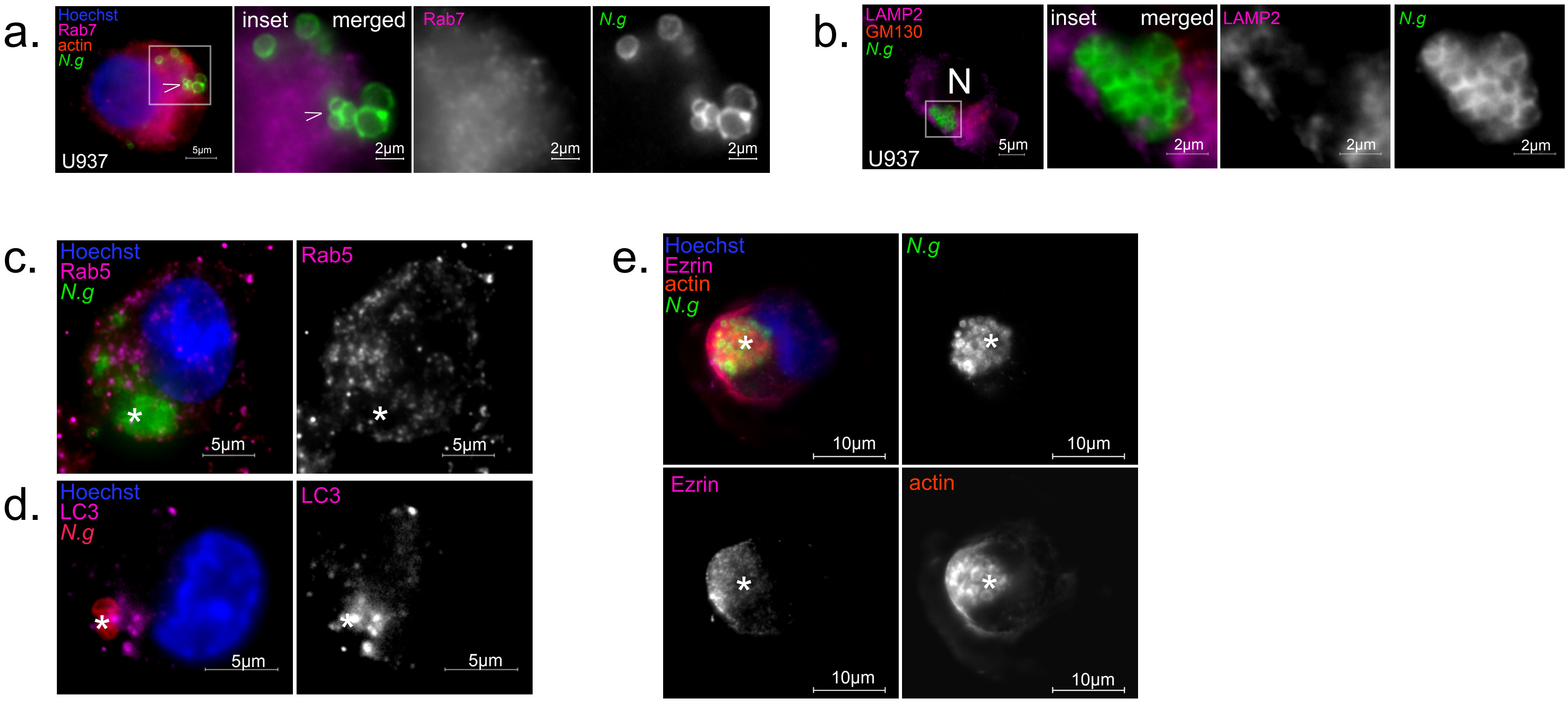

a.
Hoechst
Rab7
actin
N.g
U937
5µm
inset
merged
2µm
Rab7
2µm
N.g
2µm
b.
LAMP2
GM130
N.g
N
U937
5µm
inset
merged
2µm
LAMP2
2µm
N.g
2µm
c.
Hoechst
Rab5
N.g
5µm
Rab5
5µm
d.
Hoechst
LC3
N.g
5µm
LC3
5µm
e.
Hoechst
Ezrin
actin
N.g
10µm
N.g
10µm
Ezrin
10µm
actin
10µm

Figure S7.

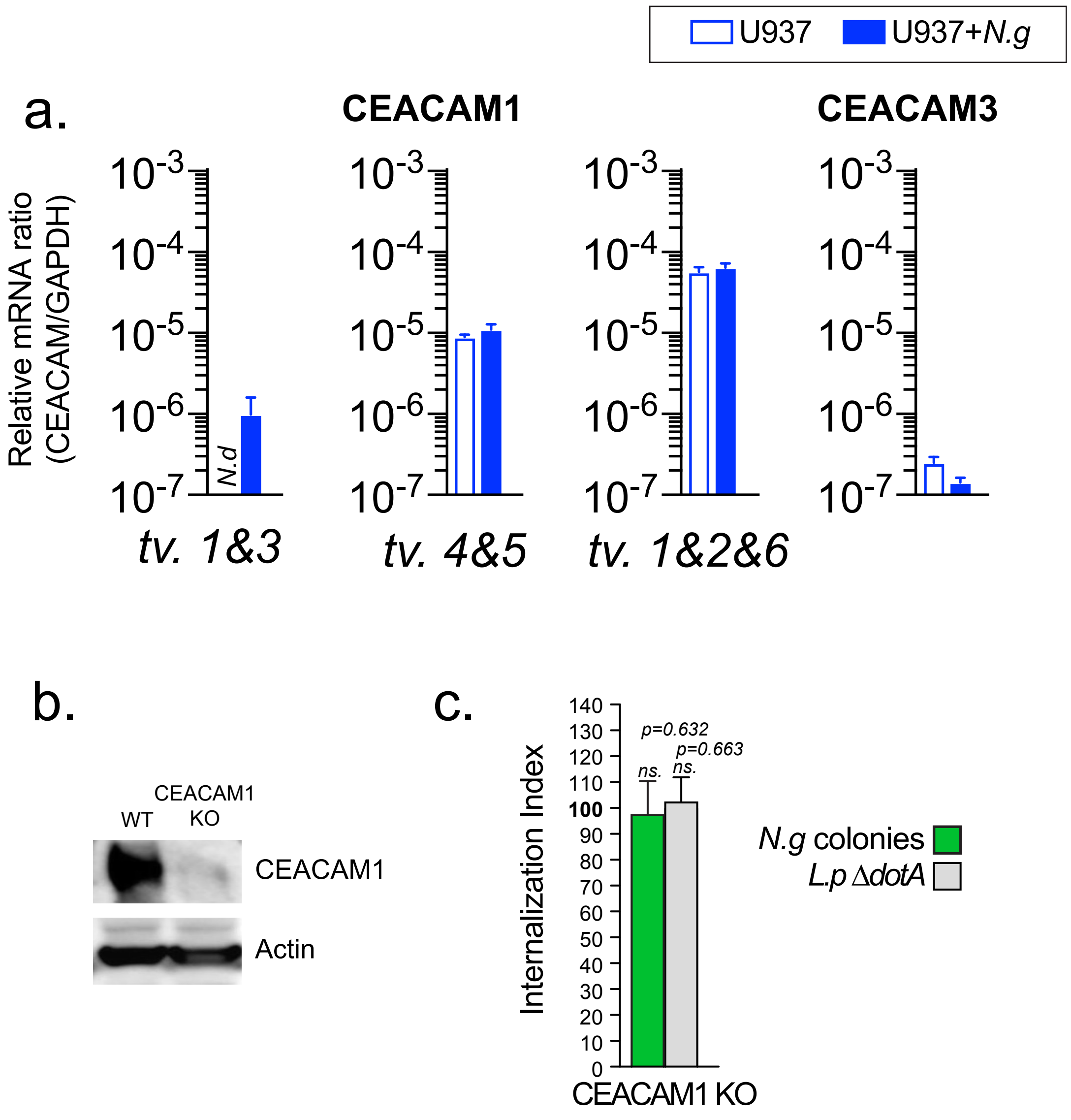
U937
U937+N.g
a.
CEACAM1
CEACAM3
CEACAM5
CEACAM6
Relative mRNA ratio (CEACAM/GAPDH)
10-3
10-4
10-5
10-6
10-7
N.d
tv. 1&3
tv. 4&5
tv. 1&2&6
tv. 1
tv. 2&3
b.
CEACAM1
WT
KO
CEACAM1
Actin
c.
Internalization Index
p=0.632
p=0.663
ns.
ns.
N.g colonies
L.p ΔdotA
CEACAM1 KO

**Supplemental Table 1. Primers and probes used for Q-PCR.** Forward (F) and Reverse (R) primers and corresponding Universal Library Probes (Roche) used for Q-PCR analysis of human formins and *ceacam* genes expression are presented. When possible, individual transcription variants (tv) were analyzed.

**Supplemental Table 2. Guide RNAs used for U937 knockout (KO) lines.** Guide RNAs used for formins, ACTR2 and CEACAM1 U937 KO lines are presented. The guide RNAs were cloned into lentiGuide-Puro (Addgene plasmid 52963).

**Supplemental Table 1**

| Name | Sequence (5’→ 3’) | Probe Number |
|---|---|---|
| F_CEACAM1_tv345 | tacctgccacgccaataact | 71 |
| R_CEACAM1_tv3 | tgggtaacatggtgagattctg | 71 |
| R_CEACAM1_tv45 | aggccattttcttgtggtagag | 71 |
| F_CEACAM1_tv126 | cccatcatgctgaacgtaaa | 57 |
| R_CEACAM1_tv126 | agggccactactccaatcac | 57 |
| F_CEACAM3 | cacagtagccctgactacagca | 1 |
| R_CEACAM3 | gtctctgctgcctgctcttt | 1 |
| F_CEACAM5_ tv1 | catagtcaagagcatcacagtctct | 2 |
| R_CEACAM5_tv1 | tcatgatgccgacagtgg | 2 |
| F_CEACAM5_tv23 | tcaagagcatcacagtctctgc | 2 |
| R_CEACAM5_tv23 | atcatgatgccgacagtgg | 2 |
| F_CEACAM6 | cggcatcacgattggagt | 21 |
| R_CEACAM6 | gaaaatacaccagggctgcta | 21 |
| F_ DAAM1_tv12 | ggagctacaagttggcctga | 46 |
| R_ DAAM1_tv12 | tccttctctaaagccagcaga | 46 |
| F_ DAAM2_tv12 | aagagctcccactgcaagac | 60 |
| R_ DAAM2_tv12 | ctttagaggcctcacccaca | 60 |
| F_ DIAPH1_tv123 | tctgcccagaccaagacttc | 39 |
| R_ DIAPH1_tv123 | tttttcttttgcacagatttctttt | 39 |
| F_ DIAPH2_tv1256 | ccaccaaacgtgagatggtt | 59 |
| R_ DIAPH2_tv1256 | tttttcagcccaccagattt | 59 |
| F_ DIAPH3_tv1235 | ttcatgcaagcaataaaggaga | 6 |
| R_ DIAPH3_tv1235 | ttctttagctattctgacacgttttt | 6 |
| F_ FMN1_tv13 | gtacccaaagccgacttgc | 1 |
| R_ FMN1_tv13 | cgatgacctatccccttgc | 1 |
| F_ FMN2 | aaaccagccacgaacactct | 6 |
| R_ FMN2 | ggtggagatgggatgttacag | 6 |

| | | |
|---|---|---|
| F_ FMNL1 | ctttgcccagtgctctgtc | 1 |
| R_ FMNL1 | tggacccttgctgaggtct | 1 |
| F_ FMNL2 | cacaacgtgcctttgaagc | 21 |
| R_ FMNL2 | ggaggtaggttcatagcattcag | 21 |
| F_ FMNL3 | tggcatataccacccatctct | 30 |
| R_ FMNL3 | gttcaagggtcccctactcc | 30 |
| F_ INF2_tv12 | gaggtctttgcctccctgtt | 32 |
| R_ INF2_tv12 | gacaggagctgggcagac | 32 |
| F_ FHOD1_tv12 | gctccctcttctcactgaagc | 79 |
| R_ FHOD1_tv12 | acaaattcaggcaccaggtc | 79 |
| F_ FHOD3_tv123 | caagacatggatttcactgacc | 52 |
| R_ FHOD3_tv123 | gacccaggtccacatctagg | 52 |
| F_ GRID2IP | ctgctgacctatgaggagca | 2 |
| R_ GRID2IP | cgtgtagcccaggaagca | 2 |

**Supplemental Table 2**

| GENE NAME | LIBRARY NUMBER | SEQUENCE |
|---|---|---|
| FMNL1 | HGLibA_17692 | GAGGGCCAACTCGCCCACCA |
| FMNL1 | HGLibA_17693 | GTGCTCCAGCACAGCGTTCT |
| FMNL1 | HGLibA_17694 | GCAGGGAGCGCACGTCCGCC |
| FMNL2 | HGLibA_17695 | ATCATACTGCCGCAGTAACC |
| FMNL2 | HGLibA_17696 | CTTTCTGCAGGAACGATTCC |
| FMNL2 | HGLibA_17697 | TCTTTGAGAACTAACCACAT |
| FMNL3 | HGLibA_17698 | CGGTGGAGGACATGAACTTC |
| FMNL3 | HGLibA_17699 | CAGAGCCATCATGAACTATC |
| FMNL3 | HGLibA_17700 | CCCAGGTATAGCACTCTCCC |
| DIAPH2 | HGLibA_13219 | GAACCGGGCCGCCAATGAAG |
| DIAPH2 | HGLibA_13220 | CCGCGCTCCGCTTGTTGCTC |
| DIAPH2 | HGLibA_13221 | CTTTAACCAGCAATCCGGTC |
| DAAM1 | HGLibA_12258 | ATCAACAATACCTCGATAGA |
| DAAM1 | HGLibA_12259 | CTCACCGGCTCATCTCAAAA |
| DAAM1 | HGLibA_12260 | CGTGTTTATGTTCCTCCAAT |
| FHOD1 | HGLibA_17491 | CTCTCCTCAGTCCCGCTTGG |
| FHOD1 | HGLibA_17492 | TACCAGAGCTACATCCTTAG |
| FHOD1 | HGLibA_17493 | GACCTCTAAGGATGTAGCTC |
| ACTR2 | HGLibA_00655 | TATAACTAGATATCTTATCA |
| ACTR2 | HGLibA_00656 | TCATTCCAGTTTGTGAAGTG |
| ACTR2 | HGLibA_00657 | CACATTTGCCCAGTATATGA |
| CEACAM1 | HGLibA_08922 | CACGCCAATAACTCAGTCAC |
| CEACAM1 | HGLibA_08923 | CATGCCATTCAATGTTGCAG |
| CEACAM1 | HGLibA_08924 | AGAGCTCTTGTGTGCTTTGC |
